# Force-responsive symmetric cell divisions orient stomata along global tissue axes

**DOI:** 10.1101/2025.11.19.689402

**Authors:** K.S. Hartman, B.Y. Lopez, J.H. Gonzalez, M.E. Goetz, A. Cleveland, A. Muroyama

## Abstract

Stomata, microscopic pores that regulate gas exchange in plants, are patterned according to conserved pathways that regulate their physiology. Here, we identify a new mode of stomatal patterning that depends on previously unrecognized regulation of the final symmetric cell division that creates paired guard cells. Symmetric cell divisions are aligned by tensile stress at both the subcellular and supracellular scales, creating a globally polarized stomatal field that tracks the major axes of tissue growth. By identifying KATANIN as a critical regulator of symmetric division orientation, we show that stress-based division orientation is required to prepattern stomatal morphology and pore creation. We find that expansion of neighboring cells non-autonomously controls symmetric division orientation, linking stomatal alignment to overall leaf shape. Finally, we show that polarized stomatal fields are widespread across plant genera and their species-specific alignment patterns are consistent with the force-based mechanism we identify in *Arabidopsis*. This force-responsive pathway provides a unifying model that explains long-standing observations of stomatal organization across species.

## Introduction

Plant morphogenesis relies on oriented cell divisions to generate cellular diversity and control tissue architecture. These divisions can be classified as either symmetric or asymmetric depending on whether they generate daughter cells with equivalent or distinct identities, respectively. Precise spatiotemporal control of both asymmetric cell division (ACD) and symmetric cell division (SCD) is essential to establish the stereotyped cellular patterns that define and regulate the physiology of various plant organs(1, 2). Hormone gradients(3–5), niche-derived soluble signals(6), mobile transcription factors(7, 8), and cell-autonomous patterning proteins(9, 10) have all been shown to regulate proliferative potential and fate acquisition depending on the developmental context. In addition to these well-established pathways, tissue mechanics are increasingly appreciated as critical drivers of organ formation(11, 12). For example, forces derived from organ and cell shape inform division patterns in the shoot apical meristem, where divisions within relatively homogenous cell populations of stereotyped geometry are oriented to minimize tensile stress(13). However, how mechanics instruct the development of other plant tissues remains an important and open question.

The leaf epidermis is a classic example of a tissue whose cellular pattern is established by pathways that coordinate oriented cell divisions with cell-fate switches(14). During leaf development, the stomatal lineage generates most of the cells and structures in the epidermis, including trichomes, pavement cells and stomata, microscopic pores formed between paired guard cells (GCs). In the early stages of the lineage in *Arabidopsis thaliana*, stomatal precursors, known as meristemoids, perform rounds of ACD that modulate the ratio of stomata to pavement cells in the leaf. Once the meristemoid stops dividing asymmetrically and becomes a guard mother cell (GMC), it performs a final SCD to create the paired guard cells that form a stoma.

Despite significant variation in cell and leaf morphology, conserved stomatal patterns can be observed across different species. The most well-characterized of these is the “one-cell spacing rule,” which depends on oriented ACDs and local inhibition of stomatal identity via small peptide signaling(15–20). In *Arabidopsis*, polarity-mediated ACD orientation regulates stomatal spacing by ensuring that new meristemoids are not created adjacent to existing GMCs and GCs. GMCs and GCs also secrete EPIDERMAL PATTERNING FACTORS (EPFs) that are perceived in early lineage cells via the ERECTA, ERECTA-LIKE and TOO MANY MOUTHS (TMM) leucine-rich repeat receptor-like kinases to locally suppress stomatal identity around existing stomata. Gas exchange measurements have confirmed that spacing is essential for proper stomatal physiology, highlighting the important link between the cellular pattern of an organ and its function(21).

Other stomatal patterns have been anecdotally noted within certain plant families(22, 23). We were particularly struck by the robust alignment of stomata along the proximodistal axis of grass leaves, which suggested that uncharacterized mechanisms control SCD orientation, at least in some species(24). Division plane defects in GMCs have been noted in *Zea mays*(*25*), *Oryza sativa*(*26*) and *Brachypodium distachyon*(*27*) mutants harboring loss-of-function alleles of MUTE orthologs, which specify GMC identity and subsidiary cell recruitment. However, the mechanistic basis underlying this SCD control in these or other plant species has not been systematically explored.

Here, we show that global stomatal alignment along leaf axes is more widespread than previously thought and is found across a range of eudicot species. By probing the mechanisms controlling this previously unrecognized SCD orientation in *Arabidopsis*, we discovered a mechanics-based pathway that operates at two length scales to couple cell-autonomous morphological alignment to global tissue axes. We demonstrate that SCD orientation is force-sensitive and the SCD field can be genetically modulated in predictable ways by altering the dominant directions of leaf expansion. Importantly, we find that SCD randomization disrupts stomatal pore formation, thereby identifying a new function for force-based SCDs during plant development. Finally, we provide evidence that this force-based mechanism controls stomatal patterns across species, providing a unifying model for polarized stomatal fields across land plants.

## Results

### Symmetric divisions that create stomata are aligned in the *Arabidopsis* leaf epidermis

Previous qualitative observations of small regions of fully developed leaves led to the conclusion that the SCDs that generate stomata are randomly oriented in *Arabidopsis*(*28*). We revisited this assumption by mapping the SCD landscape across the leaf epidermis, which can be derived from the placement of the cell walls separating paired GCs (Figure S1A). We confirmed that this method faithfully reports SCD orientation, even following cell expansion (Figure S1B). Annotating all SCDs across the entire abaxial epidermis of 3 days post-germination (3 dpg) cotyledons revealed that, contrary to previous assumptions, division orientation is strongly biased along the cotyledon’s proximodistal axis (Figure 1A). As stomatal distribution on each cotyledon or true leaf is unique, we developed a method to generate a representative, composite leaf to visualize the average SCD orientation across the epidermis (see Methods). Applying this method to our annotated 3 dpg cotyledon revealed a robustly polarized SCD field globally aligned along the proximodistal axis (Figure 1B).

**Figure 1:**
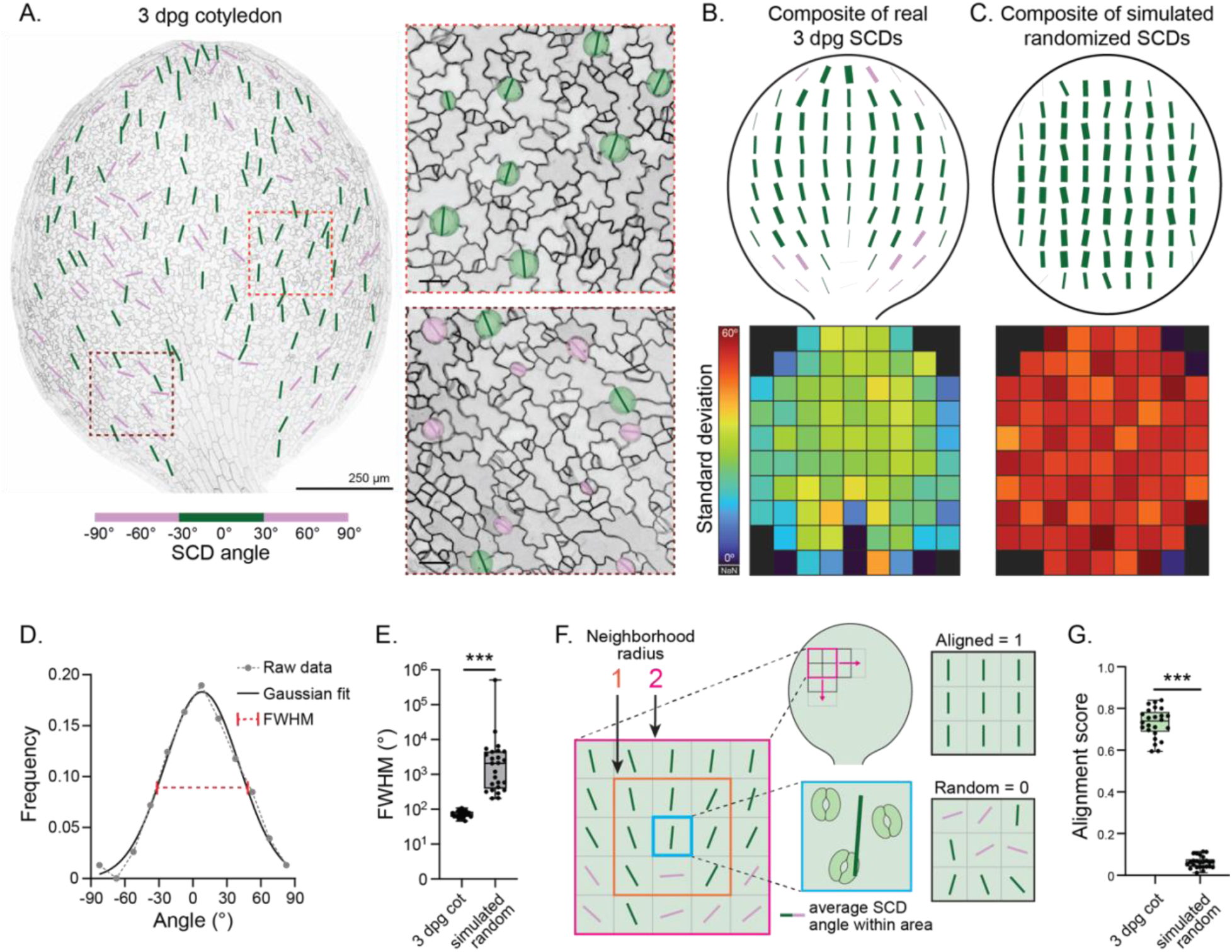
Symmetric cell divisions in guard mother cells are aligned in the *Arabidopsis* leaf epidermis. A. Representative image of a 3 dpg ML1p::mCherry-RCI2A-expressing cotyledon (left) and zoomed regions (right) with the SCDs color-coded according to their orientation relative to the proximodistal axis. Scale bars – 250 μm (left) and 25 μm (right). B. (Top) Composite 3 dpg cotyledon generated from experimentally derived data showing the average SCD orientation across the epidermis. (Bottom) Associated map showing the corresponding standard deviations for SCD orientation at each position. n = 24 cotyledons. C. (Top) Composite “cotyledon” from simulated cotyledons with randomly oriented SCDs, showing the average SCD orientation across the surface. (Bottom) Associated map showing the corresponding standard deviations for SCD orientation at each position. n = 25 cotyledons. D. The distribution of SCD angles from the example in (A) and associated Gaussian fit and full-width half max (FWHM) value. E. FWHM values of the 3 dpg and simulated, randomized cotyledons from (B) and (C), respectively. n = 24 (3 dpg) and 25 (simulated) cotyledons. F. Alignment by Fourier Transform (AFT) method for calculating local alignment among neighboring stomata. G. Alignment scores for the 3 dpg and simulated, randomized cotyledons from (B) and (C), respectively. n = 24 (3 dpg) and 25 (simulated) cotyledons. *** - p<0.001

If SCD orientation were random in individual cotyledons, with stomatal angles uniformly distributed between -90° and 90° around the proximodistal axis (0°), the average orientation of the stomatal field derived from many independent samples would be biased along the proximodistal axis, a prediction we verified by simulating cotyledons with randomized SCD orientation (Figure 1C and Figure S1C). Therefore, to confirm that SCD orientation is non-random in 3 dpg cotyledons, we developed quantitative measures to compare three parameters of stomatal alignment. (1) The between-sample variability of SCD orientation at a given position within the tissue can be assessed by calculating the standard deviation of SCD angles at defined intervals across the leaf epidermis. SCD angle was highly variable between the simulated randomized samples, as expected, while variability was low between real 3 dpg cotyledons, demonstrating consistent stomatal alignment between cotyledons (Figure 1B,C). (2) Global SCD alignment to the proximodistal axis can be measured by fitting the distribution of SCD angles within a single sample with a Gaussian function and calculating the full-width half-maximum (FWHM) value; smaller and larger FWHM values represent higher and lower alignment, respectively (Figure 1D). Stomata were consistently aligned with the proximodistal axis in 3 dpg cotyledons but poorly aligned in simulated samples (Figure 1E). (3) Local concordance of SCD orientation among neighboring stomata can be quantified using a modified version of Alignment by Fourier Transform (AFT)(29), a program developed to measure alignment of fibers within an image (Figure 1F). Again, SCDs showed strong local alignment in 3 dpg cotyledons but very poor alignment in our randomized samples (Figure 1G). Therefore, by all three metrics, stomatal alignment in 3 dpg *Arabidopsis* cotyledons is non-random and strongly biased along the proximodistal axis.

Next, we evaluated if global SCD orientation changes over development by quantifying stomatal alignment in 3 to 7 dpg cotyledons, over which time there is an ∼4-fold increase in the number of stomata per cotyledon. Stomatal alignment to the proximodistal axis and alignment between neighboring stomata both decreased from 3 to 5 dpg, although they always remained biased in the proximodistal direction (Figure S1D-F). Therefore, SCD orientation changes dynamically over cotyledon development but always remains non-random. Finally, we checked whether stomatal alignment is specific to cotyledons or is found in other stomata-containing aerial tissues. True leaves also showed global stomatal alignment biased along the proximodistal axis and, interestingly, SCDs were almost universally oriented along the proximodistal axis in sepals (Figure S1G-K). From these data, we conclude that division orientation in GMCs is actively regulated across *Arabidopsis* tissues, providing a new experimental system to explore the mechanisms controlling polarized SCDs during plant morphogenesis.

### Key regulators of polarized division and cell fate in the stomatal lineage do not control SCD orientation

To determine how GMC SCDs are oriented, we started by testing if known regulators of cell division in the stomatal lineage are required to generate the polarized stomatal field. ACDs in the *Arabidopsis* stomatal lineage are oriented cell-autonomously by the plasma membrane-associated polar proteins BREAKING OF ASYMMETRY IN THE STOMATAL LINEAGE (BASL) and members of the BREVIS-RADIX family (BRXf)(9, 30). Even though fluorescent BASL and BRXf reporters can be planar polarized along global leaf axes(31–33) and our reporters showed persistent, albeit nonpolar, expression in GMCs (Figure S2A), we found that SCD orientation was unchanged in *basl-2* and *brx-q*, a quadruple, loss-of-function mutant in four of the five BRXf genes (Figure 2A,B). The OCTOPUS-LIKE (OPL) protein family was recently shown to polarize opposite to BASL and BRXf in asymmetrically dividing cells and, intriguingly, OPL2 localized to the shared cell wall separating GCs immediately after division(34). We found that SCDs continued to be oriented along the proximodistal axis in the quadruple *opl1-1 opl2-3 opl3-1 opl4-1* mutant (*opl-q*), indicating that the polar proteins controlling ACD and division potential are not required for SCD orientation (Figure 2A,B).

**Figure 2:**
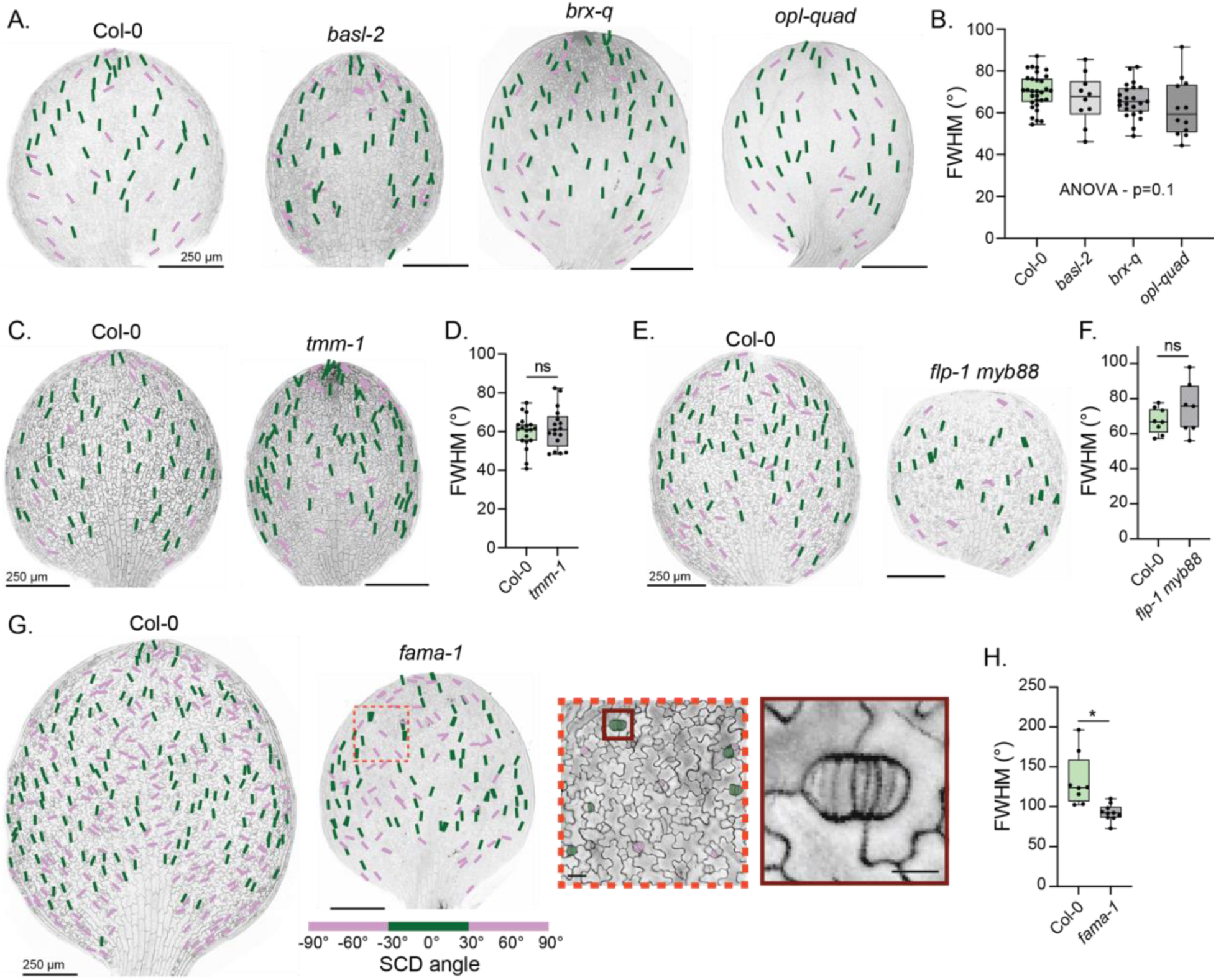
SCD orientation is not controlled by key stomatal lineage regulators. A. Representative images of 3 dpg cotyledons from the indicated genotypes (Col-0, *basl-2*, *brx-q*, *opl-quad*) with color-coded SCDs. Scale bars – 250 μm. B. FWHM values for Col-0, *basl-2*, *brx-q*, and *opl-quad*. n = 31 (Col-0), 10 (*basl-2*), 23 (*brx-q*) and 12 (*opl-quad)* cotyledons. C. Representative images of 3 dpg Col-0 and *tmm-1* cotyledons with color-coded SCDs. Scale bars – 250 μm. D. FWHM values for Col-0 and *tmm-1*. n = 19 (Col-0) and 17 (*tmm-1*) cotyledons. E. Representative images of 3 dpg Col-0 and *flp myb88* cotyledons with color-coded SCDs. Scale bars – 250 μm. F. FWHM values for Col-0 and *flp myb88*. n = 8 (Col-0) and 7 (*flp myb88*) cotyledons. G. Representative images of 5 dpg Col-0 and *fama-1* cotyledons (left) and zoomed regions showing parallel, oriented SCDs in *fama-1* (right) with color-coded SCDs. Scale bars – 250 μm (two leftmost images), 25 μm (middle image), and 10 μm (right image). H. FWHM values for Col-0 and *fama-1*. n = 8 (Col-0) and 10 (*fama-1*) cotyledons. ns – not significant, * - p<0.05

Next, we tested whether key stomatal fate regulators are essential for oriented GMC SCD. *TMM* mutants show misoriented ACDs and aberrant induction of stomatal fate, leading to the formation of stomatal clusters(15, 35). Global SCD alignment was unchanged in a *TMM* loss-of-function mutant (*tmm-1*), demonstrating that EPF signaling is dispensable for SCD orientation (Figure 2C,D). The transcription factors FOUR LIPS (FLP) and MYB88 are important regulators of the GMC to GC transition, and *FLP MYB88* mutants exhibit stomatal patterning defects, most notably paired stomata(36). Similar to our observations in *tmm-1*, we found no defects in SCD orientation in *flp-1 myb88* (Figure 2E,F). Along with our analyses of *basl* and *brx-q*, these results indicate that one-cell spacing is not required for oriented SCDs.

Finally, we checked if the transcription factor FAMA is necessary for SCD orientation. *FAMA* expression is initiated in late-stage GMCs before they undergo SCD and is required to both limit GMC divisions and establish proper guard cell identity(37). In *FAMA* mutants, GMCs continue to divide, creating “*fama* tumors” that never form stomatal pores. Surprisingly, even though *FAMA* expression coincides with the SCD and is necessary for guard cell differentiation, it is not strictly required for SCD orientation. Even though overall cotyledon growth was slightly suppressed in *fama-1*, GMC divisions continued to be aligned with the proximodistal axis in *fama-1* at a frequency comparable to that seen in wild-type cotyledons of similar size (Figure 2G,H). Taken together, our mutant analyses reveal that neither known regulators of stomatal lineage ACD nor transcription factors controlling GMC division and guard cell fate are necessary for oriented GMC SCDs.

### GMC division orientation is not controlled by a vein-secreted signal

Having found that GMC SCDs are not oriented by canonical regulators of stomatal patterning and fate, we tested the contribution of other high-priority candidate pathways. Stomatal formation is coordinated with the development of neighboring tissues via mobile signals(38). Because the SCD field closely mirrored the organization of the underlying vasculature and a tight relationship between stomatal distribution and veins has been noted in some species(39–41), including grasses(23, 42), we tested whether division orientation in GMCs depended on proper vein specification. To do this, we measured SCD orientation in two mutants, *tir1-1 afb2-3 afb3-4*(*43*) and *scarface* (*scf-9*)(44), that have defects in vein development. SCD orientation across the leaf epidermis was unaffected in both of these mutant backgrounds, leading us to conclude that GMC division orientation does not depend on the pre-patterned secretion of a vein-derived mobile signal (Figure S2B-E).

### Auxin does not drive polarized SCD orientation

Auxin gradients control cell division patterns in developing plant tissues(45, 46). Within the stomatal lineage, auxin activity fluctuates dynamically during differentiation, and mutants defective in auxin signaling exhibit aberrant stomatal patterning, including asymmetric division defects(47, 48). However, auxin’s role in the final SCD has not been characterized. We tested whether auxin-mediated signaling controls GMC division orientation by measuring stomatal alignment in seedlings grown on plates containing the synthetic auxin analog indole-3-acetic acid (IAA) or *N*-1-naphthylphthalamic acid (NPA), an inhibitor of the PIN auxin efflux transporters(49). As expected, both IAA and NPA treatments suppressed cotyledon development in a dose-dependent manner (Figure S2F,H). However, despite changes in overall cotyledon size, neither treatment negatively disrupted GMC division orientation in 3 dpg seedlings (Figure S2F-I). We noted a slight increase in stomatal alignment in our seedlings exposed to 1 μM IAA, but as the FWHM values were similar to those observed in Col-0 seedlings with similar stomatal counts, we believe that this change reflects an IAA-induced delay in cotyledon development rather than a specific effect on SCD orientation. Together with the observation that SCD orientation is not altered in *tir1-1 afb2-3 afb3-4* (Figure S2D,E), in which auxin signaling is also disrupted, we conclude that auxin is not a critical regulator of polarized SCDs in the stomatal lineage.

### GMC morphology predicts division orientation

Several models have been developed to describe how cell division can follow morphological or mechanical cues in the absence of other extrinsic or intrinsic signals. The Besson-Dumais rule, which states that symmetrically dividing plant cells probabilistically select from a range of possible division orientations that minimize the new cell wall’s surface area, built on Errera’s rule and explains the division patterns observed in some but not all plant species and tissues(50). The rules describing SCD orientation were further generalized and refined by Louveaux et al., who determined that SCD orientation in the *Arabidopsis* shoot apical meristem was better explained by a mechanical rule where cells divided along the axis of maximal tensile stress(13).

Anecdotal descriptions of GMC behavior suggested that they may divide along their long axes(51, 52). To systematically examine whether GMC morphology robustly predicts SCD orientation, we performed time-lapse imaging in seedlings expressing an epidermal-specific plasma membrane marker (ML1p::mCherry-RCI2A) that allowed us to monitor GMC growth and associated division patterns. In our time-lapse movies, GMC morphology and division orientation were strongly correlated, with the newly created cell wall closely matching the GMC’s long axis (R^2^ = 0.76) (Figure 3A,B), suggesting that cell shape might be an important determinant of SCD orientation in the stomatal lineage.

**Figure 3:**
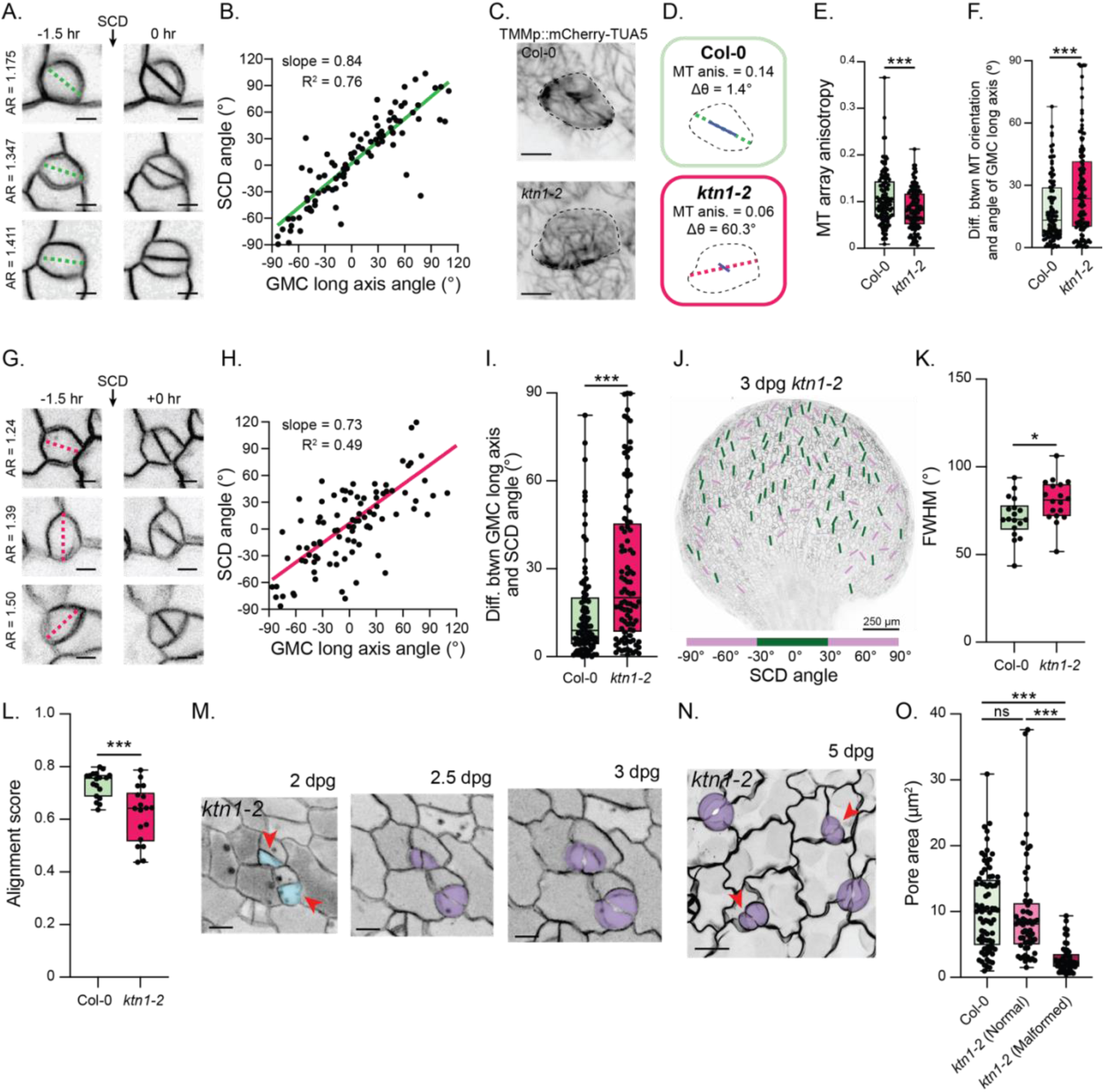
SCD orientation follows GMC morphology and requires KATANIN. A. Representative images of GMC divisions from 3 dpg ML1p::mCherry-RCI2A-expressing cotyledons. The dashed green line shows the calculated longest axis before division. Scale bars – 5 μm. B. Comparison of the angle of the GMC long axis immediately before mitosis to the associated SCD angle (green line – linear fit). n = 91 cells. C. Representative images of microtubule organization in TMMp::mCherry-TUA5-expressing GMCs in Col-0 and *ktn1-2* cotyledons. Scale bars – 5 μm. D. Simplified representation of the GMCs in (C) showing the cell outline (thin dashed line), the GMC long axis (dashed green or magenta line), and the major direction of microtubule alignment (solid blue line). E. Anisotropy of the microtubule network in Col-0 and *ktn1-2* GMCs. n = 125 (Col-0) and 124 (*ktn1-2*) cells. F. Differences between the orientation of the microtubule array and the GMC long axis. n = 103 (Col-0) and 124 cells (*ktn1-2*) cells. G. Representative images of abnormally oriented SCDs in *ktn1-2* (dashed magenta line – calculated longest axis). Scale bars – 5 μm. H. Comparison of the angle of the GMC long axis immediately before mitosis to the associated SCD angle in *ktn1-2* (magenta line – linear fit). n = 96 cells. I. Differences between the GMC long axis angle and the angle of the SCD. n = 91 (Col-0) and 96 cells (*ktn1-2*) cells. J. Representative image of 3 dpg *ktn1-2* cotyledon with color-coded SCDs. Scale bar – 250 μm. K. FWHM values for Col-0 and *ktn1-2* 3 dpg cotyledons. n = 18 cotyledons each. L. Alignment scores for the Col-0 and *ktn1-2* cotyledons in (D). n = 18 cotyledons each. M. Representative example of misoriented *ktn1-2* SCDs tracked over 24 hours. Scale bars – 10 μm. N. Representative image of a 5 dpg ML1p::mCherry-RCI2A-expressing *ktn1-2* cotyledon with malformed stomata (red arrows). Scale bar – 25 μm. O. Pore areas in 5 dpg Col-0 and *ktn1-2* cotyledons. n = 76 (Col-0), 57 (normal, *ktn1-2*), and 50 (malformed, *ktn1-2*) pores. ns – not significant, * - p<0.05, *** - p<0.001

Wild-type GMCs expand before undergoing SCD, approaching circularity before mitotic entry (median circularity = 0.907). This raised the possibility that the nearly isotropic GMC shape would introduce some degree of stochasticity in division plane selection, similar to what has been seen in other cases(53). Indeed, SCD angles in Col-0 better matched the GMC long axis in more anisotropic GMCs, and SCDs in highly isotropic GMCs deviated most significantly from the GMC long axis (Figure S3A,B). Therefore, to test whether increasing GMC anisotropy is sufficient to increase the predictive power of GMC morphology, we identified a mutant that affects GMC shape. Cortical microtubules control cell shape across cell types(54), although no work, to our knowledge, has systematically quantified GMC morphology in seedlings harboring mutations in microtubule-associated proteins. In our previous study(55), we noted that epidermal cells were misshapen in the *trm678* mutant, which lacks three TON1-RECRUITING MOTIF (TRM) proteins(56). Having found that GMCs were significantly more anisotropic in *trm678* than wild-type (Figure S3C), we performed time-lapse imaging in *trm678* ML1p::mCherry-RCI2A seedlings and found that increasing GMC anisotropy significantly improved concordance between the GMC long axis and the SCD angle (R^2^ = 0.95) (Figure S3D-F). From these data, we conclude that GMCs divide along their long axes and that their near circularity before mitotic onset in Col-0 introduces some stochasticity in division plane selection during wild-type GMC divisions.

### KTN1 controls microtubule organization and division orientation in GMCs

In other plant tissues, cells that divide along their long axes do so because they follow the direction of greatest tensile stress(13). No work has yet identified a role for mechanical forces in controlling division orientation during stomatal patterning, but our time-lapse data suggest that force-based cues might be the dominant regulator of division orientation in *Arabidopsis* GMCs.

The magnitude and direction of the stresses within individual plant cells are difficult to experimentally determine, but cortical microtubules are generally considered to be force-responsive in plants and will typically align parallel to the direction of greatest stress(57). Using a stomatal lineage-specific microtubule reporter line (TMMp::mCherry-TUA5), we quantified microtubule anisotropy and alignment on the periclinal surface of GMCs and found that microtubules were well-aligned along the GMC long axis (Figure 3C-F). Because microtubule ordering along the axis of greatest stress is dependent on the microtubule-severing protein KATANIN1 (KTN1) in the shoot apical meristem(58), we quantified microtubule organization in *ktn1* GMCs by introgressing the TMMp::mCherry-TUA5 reporter into *ktn1-2*. Microtubule network anisotropy and alignment along the GMC long axis were both decreased upon *KTN1* loss (Figure 3C-F), indicating that the GMC long axis is indeed its axis of maximum stress.

*ktn1* GMCs are slightly more anisotropic than those in Col-0 (Figure S3I), so we expected to find improved concordance between SCD orientation and the GMC long axis if divisions were strictly based on cell morphology, as we saw in *trm678*. If, instead, GMCs are dividing along the axis of greatest stress, we would predict that SCD orientation would be perturbed in *ktn1*-*2.* To distinguish between these two possibilities, we generated a *ktn1-2* ML1p::mCherry-RCI2A line and performed time-lapse imaging to monitor GMC behavior in 3 dpg cotyledons. We found increased randomization of SCD orientation in *ktn1-2*, with many SCDs that bisected the GMC along a shorter path, demonstrating that KTN1 is required to orient GMC divisions in the *Arabidopsis* stomatal lineage and implicating force as a key regulator of division orientation (Figure 3G-I). Accordingly, global SCD alignment along the proximodistal axis was significantly decreased in *ktn1-2*, as was the alignment between neighboring stomata (Figure 3J-L). Notably, the striking variability in local alignment between *ktn1-2* cotyledons indicated that the decreased FWHM values in *ktn1-2* are not due to a lateral reorientation of SCDs but rather a significant increase in SCD randomization (Figure 3L). Taken together, these data are consistent with a model where GMC divisions are oriented along the axis of greatest stress, which both 1) informs and is read out by the major axis of the cortical microtubule array and 2) coincides with the long axis in Col-0 GMCs.

The SCD randomization in *ktn1-2* presented an opportunity to test whether defects in SCD orientation have phenotypic consequences on stomatal formation. By monitoring epidermal development from 2 to 3 dpg in *ktn1-2* cotyledons, we could measure SCD orientation relative to the GMC long axis and follow associated guard cell maturation and pore formation within single cells (Figure 3M). All SCDs created volumetrically equal daughters, but guard cells that were created from SCDs that failed to align along the GMC long axis exhibited striking morphological defects during guard cell maturation. By 5 dpg, the *ktn1-2* epidermis contained two distinct classes of stomata: those with kidney-shaped guard cells that were morphologically indistinguishable from Col-0 stomata and those that were misshapen, with increased lateral expansion and reduced expansion along the shared wall (Figure 3N). In both cases, microtubules were organized in the characteristic radial arrays that define interphase microtubules in guard cells, indicating that malformed stomata in *ktn1-2* are not caused by gross differences in microtubule organization post-division (Figure S3J). Notably, pore area was significantly decreased in malformed stomata in *ktn1-2* but was unaffected in normal *ktn1-2* stomata, which had identical pore areas as those formed in Col-0 (Figure 3O, S3K). These results identify a previously unrecognized function for oriented SCD in controlling guard cell morphology and pore formation.

### Developmentally regulated growth patterns globally align GMC SCDs

Our time-lapse data demonstrated that GMC morphology cell autonomously predicts SCD orientation of any individual cell but does not, on its own, explain why SCDs are oriented in a polarized field across the tissue. Indeed, while concordance between the GMC long axis and division plane is increased in *trm678*, global alignment is actually reduced relative to Col-0 cotyledons (Figure S3G,H). While analyzing GMC behavior and SCD orientation in wild-type and stomatal patterning mutants, we made several observations that hinted that anisotropic GMC expansion might align neighboring stomata to generate the polarized SCD field across the epidermis.

(1) In Col-0 cotyledons, we observed subtle regional differences in the direction of global alignment across the epidermis. Notably, SCDs near the petiole fanned outward along the leaf margins and appeared well-aligned with each other even though they were less globally aligned along the proximodistal axis (Figure 1A). Our analysis of alignment between neighboring stomata confirmed that distant stomata are significantly less aligned than ones in close proximity (Figure S4A). Differences in local stomatal alignment were magnified in the serrations around the edges of true leaves, where SCDs were oriented towards the serration tip instead of the proximodistal axis (Figure S4B).
(2) Stomatal alignment differed significantly between the 3 dpg abaxial and adaxial surfaces (Figure S4C), which modeling suggests principally grow in different directions to generate the cup-shaped cotyledon(59, 60).
(2) Stomatal clusters can arise from GMC divisions that occur at the same time (synchronous) or at distinct developmental stages (asynchronous). Closer examination of the paired stomata in *basl-2* and *flp-1 myb88* revealed that synchronous pairs were well-aligned with each other while asynchronous ones were significantly less well-aligned (Figure S4D,E). This observation suggested that the GMC position alone is not sufficient to predict GMC orientation. Instead, SCD orientation is dependent on the interaction between GMC position and developmental stage.
(4) While screening for mutants with altered SCD orientation, we identified two, *kan1-11 kan2-5* and *rot3-1*, where leaf shape and stomatal alignment were both changed. Mutants harboring loss-of-function alleles in the transcription factors *KANADI1* (*KAN1*) and *KANADI2* (*KAN2*) exhibit an adaxialized abaxial leaf surface and significantly elongated true leaves(61), while mutations in *ROTUNDIFOLIA3* (*ROT3*) result in squatter leaves(62). Cotyledon morphology is not perturbed in either genetic background, but differences in true leaf shape were readily observed by 10 dpg and can be traced, in part, to changes in pavement cell expansion. We found that stomatal alignment along the proximodistal axis was decreased in the squat *rot3-1* leaves and that stomata were hyper-aligned in the elongated 10 dpg *kan1-11 kan2-5* true leaves (Figure S5), revealing a link between cellular expansion and SCD orientation.

To directly measure the evolution of GMC shape and stomatal formation, we quantified GMC orientation, expansion and eventual SCD orientation in 623 GMCs for 24 hours from 2 dpg to 3 dpg (Figure 4A) (see Methods for detailed explanation). Briefly, we sorted all tracked GMCs into three groups corresponding to how well their long axes at time = 0 hrs (2 dpg) aligned with the average SCD angle at that position at time = 24 hrs (3 dpg). We then used hierarchical clustering as an unbiased way to classify the average GMC growth and division patterns. The resulting nine clusters represented the nine possible trajectories a cell could follow from initial to final alignment (Figure 4B,C). We found that the overwhelming majority (96%) of GMCs that were initially oriented along the predicted angle at that position maintained that orientation during their expansion phase and produced aligned SCDs. In contrast, half (50%) of GMCs with initially unaligned long axes reoriented through cell expansion to become more aligned by the time they underwent SCD.

**Figure 4:**
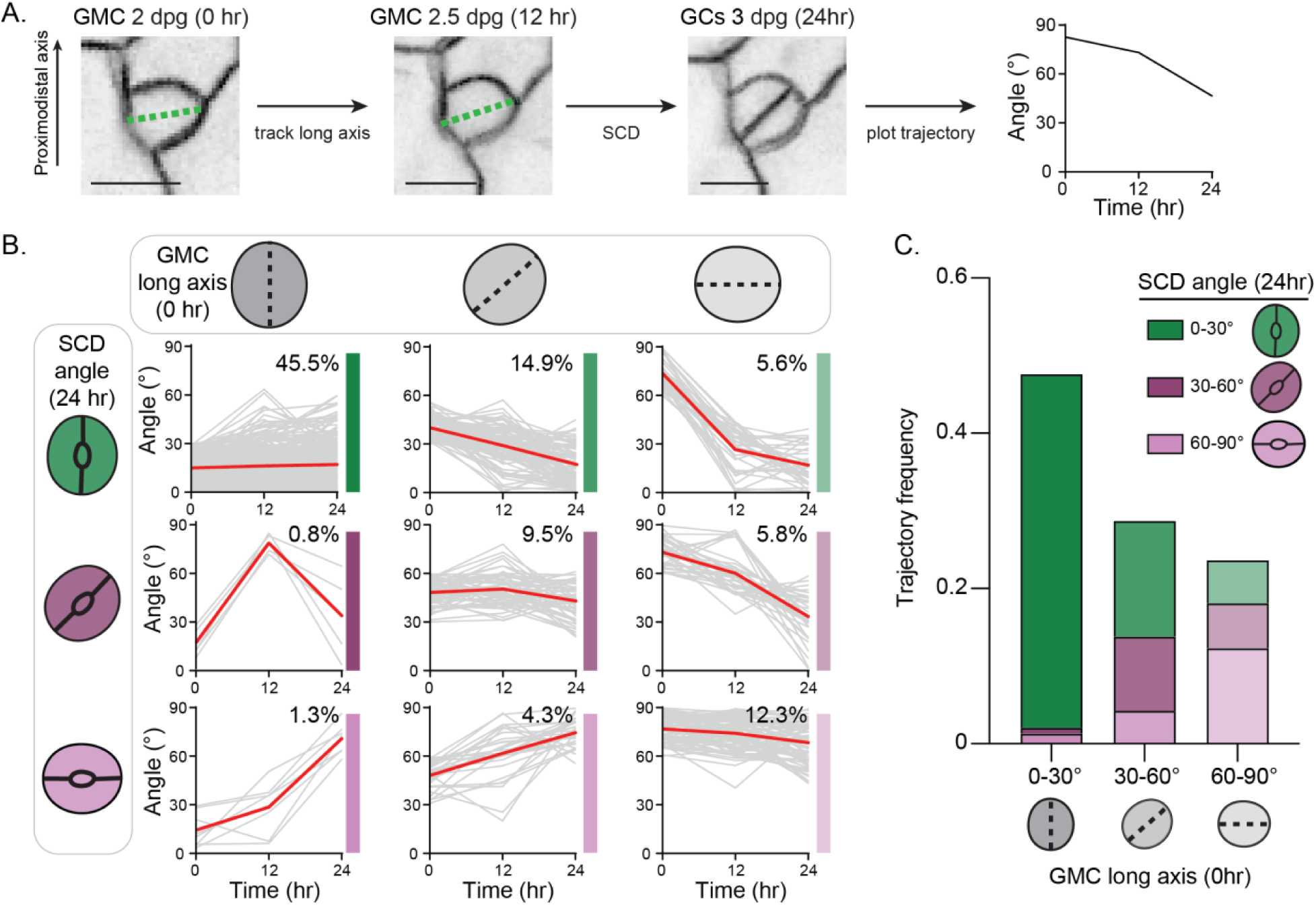
Directional GMC expansion primes globally aligned SCDs. A. Representative example of GMC tracking over 24 hours from 3 to 4 dpg. Scale bars – 5 μm. B. Major patterns of directional cell expansion and SCD orientation following hierarchical clustering. The red lines indicate the average growth and SCD orientation for a given GMC class. The percent of total GMCs that fell within each class are shown in the upper right corners and the colored bars to the right correspond to the GMC behavioral categories shown in (C). n = 623 cells. C. GMC behavioral categories based on hierarchical clustering.

### Altering cell growth through cell ablation is sufficient to reorient SCD orientation

Because our tracking data supported a model where GMC expansion pre-patterns the oriented stomatal field, we next wanted to test whether changing the direction of GMC growth would be sufficient to reorient SCD orientation. The magnitude and direction of cellular growth are controlled not only by a cell’s internal turgor pressure and its own cell wall but are also influenced by the mechanical environment dictated by the surrounding cells(63, 64). Previous work showed that small ablations in the leaf epidermis reorient tensile stress in the neighboring cells surrounding the ablation site, triggering microtubule reorganization within hours(65). Hypothesizing that the ablation and subsequent microtubule reorganization would locally affect cell expansion, we generated small wounds on one side of the cotyledon and assayed whether this physical perturbation was sufficient to reorient SCD. First, we measured SCD alignment relative to the ablation and an identically shaped non-ablated control region on the contralateral side of the same cotyledon (Figure 5A,B). As expected, stomata were not aligned relative to the ablation site upon initial wounding (Figure 5C,D). In contrast, after 24 hours, we found that SCDs significantly reoriented towards the ablation center but remained randomly oriented on the control side (Figure 5E,F).

**Figure 5:**
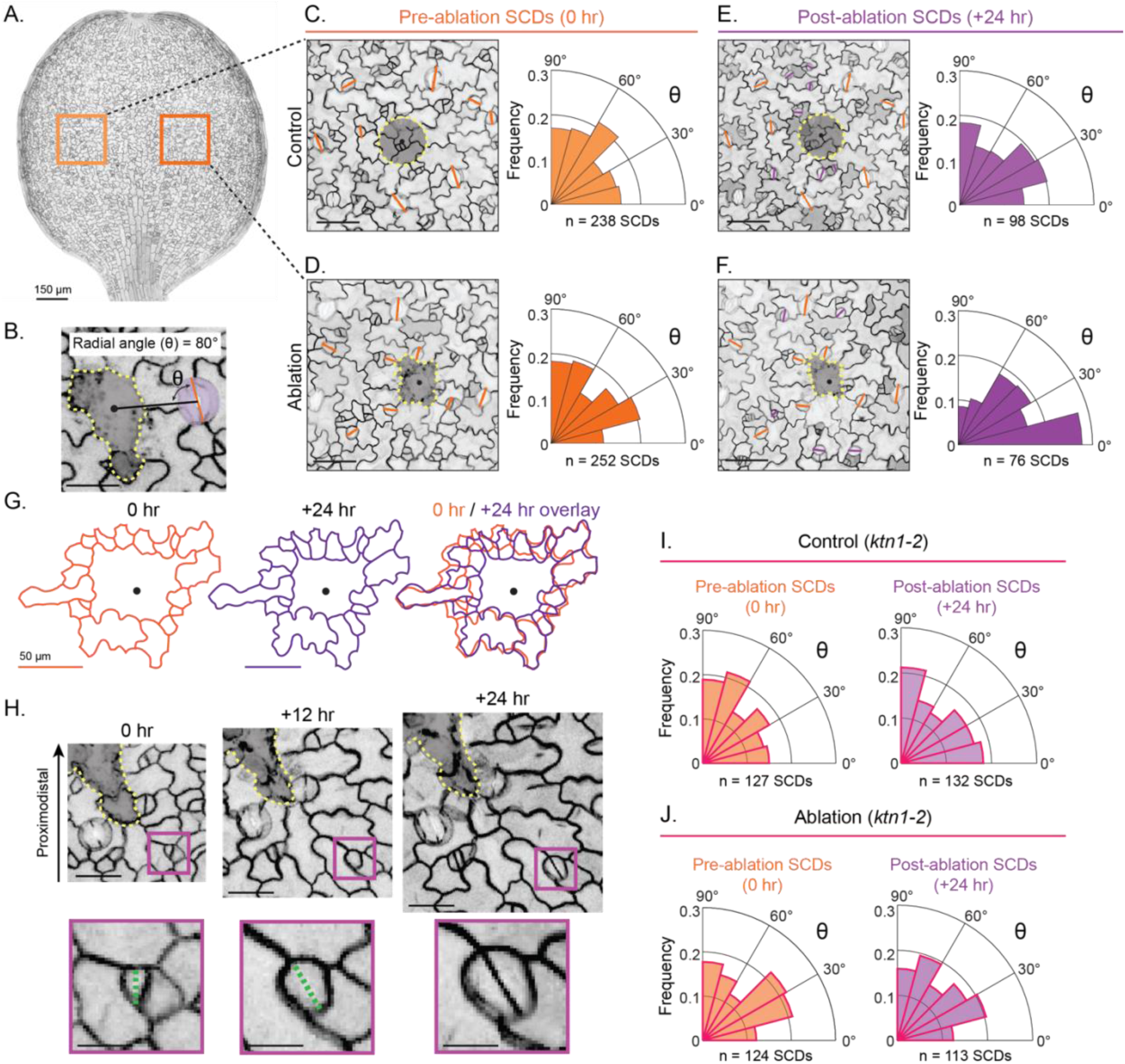
Ablations are sufficient to reorient neighboring SCDs. A. Representative image of a 3 dpg cotyledon where the SCDs around either an ablation or contralateral control site were monitored. Scale bar – 150 μm. B. Illustration of how the radial angle between the ablation site (dashed yellow line) and the SCD (solid orange line) was calculated. Scale bar – 25 μm. C-F. SCD tracking around control (C, E) and ablation sites (D, F). (Left) Representative images of the tracked areas. (Right) Polar histograms of the radial angles (θ). Scale bars – 50 μm. G. Representative example of the cellular front surrounding the ablation site at 0 and 24 hours. Scale bars – 150 μm. n = 16 cotyledons. H. (Top) Representative images of a GMC reorienting its long axis towards the ablation site (dashed yellow line) and the subsequent SCD. Scale bars – 25 μm. (Bottom) Zoomed image of the indicated GMC and its long axis (dashed green line). Scale bars – 10 μm. I-J. Polar histograms of the radial angles (θ) around control (I) and ablation sites (J) in *ktn1-2* cotyledons. n = 26 cotyledons.

To analyze how the ablations caused SCD reorientation, we monitored GMC behavior in the 24 hours following ablation, similar to our analysis of GMC growth and division in unperturbed 2-3 dpg cotyledons. We found that ablations shifted the major direction of cellular growth, reorienting expansion towards the wound end (Figure 5G,H). Finally, to confirm that this reorientation depended on changes in mechanical force, we tested whether KTN1 was required for ablation-induced SCD reorientation and found that SCDs did not reorient towards the wound in *ktn1-2* (Figure 5I,J). From these experiments, we conclude that SCDs are force-responsive and that altering the major direction of cell expansion is sufficient to reorient SCD.

### Genetically induced changes to cellular growth are sufficient to reorient the global SCD pattern

Our ablation data demonstrated that SCDs can be locally reoriented by changing the major direction of tensile stress but were not sufficient to determine whether this mechanism could generate a global pattern during leaf development. To comporehensively test this hypothesis, we created a series of *Arabidopsis* lines where the magnitude and principal direction of cell expansion could be titrated within distinct cellular populations during leaf development. First, we generated plants overexpressing *LONGIFOLIA1* (*LNG1*, also known as *TRM2*), as 35Sp::LNG1 plants were previously shown to have dramatically elongated pavement cells and leaves(66, 67). Stomatal lineage phenotypes in LNG1-overexpressing lines were not previously characterized.

We overexpressed LNG1 using three different promoters: 1) the ubiquitously expressed and constitutively active *UBIQUITIN10* promoter (UBQ10p), 2) the stomatal lineage-specific *TMM* promoter (TMMp), and 3) the early stomatal lineage-specific *SPEECHLESS* (SPCHp) promoter. All lines were generated in an ML1p::mCherry-RCI2A background, allowing us to track development. All three genotypes developed elongated cotyledons and true leaves but the degree of anisotropy varied based on the promoter; UBQ10p::LNG1 cotyledons were the most elongated, followed by TMMp::LNG1 and SPCHp::LNG1 cotyledons, respectively (Figure 6A, Figure S6A).

**Figure 6:**
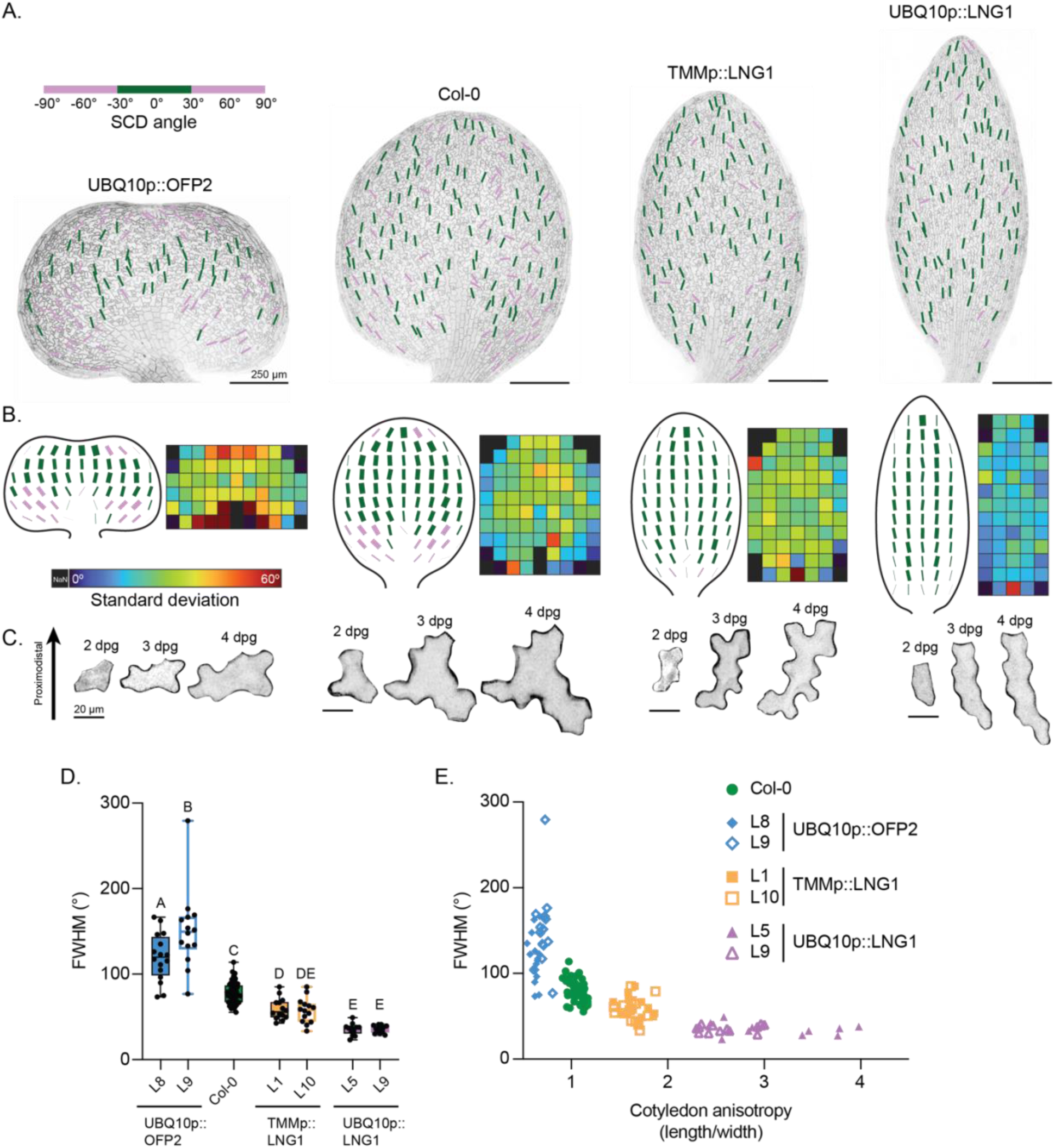
Altered expansion in neighbor cells is sufficient to reorient the polarized SCD field. A. Representative images of 3 dpg cotyledons from the indicated genotypes (UBQ10p::OFP2 L8, Col-0, TMMp::LNG1 L1, UBQ10p::LNG1 L9) with color-coded SCDs. Scale bars – 250 μm. B. (Left) Composite 3 dpg cotyledons showing the average SCD orientation by position for each genotype shown in (A). (Right) Associated maps showing the corresponding standard deviations for SCD orientation at each position. n = 16 (UBQ10p::OFP2 L8), 55 (Col-0), 14 (TMMp::LNG1 L1) and 13 (UBQ10p::LNG1 L9) cotyledons. C. Representative image series of typical pavement cell expansion for each of the genotypes in (A), oriented with respect to the cotyledon proximodistal axis. Scale bars – 20 μm. D. FWHM values from two independent 3 dpg transgenic lines for each indicated genotype. Same n values as (B), and for additional lines, n = 14 (UBQ10p::OFP2 L9), 14 (TMMp::LNG1 L10) and 13 (UBQ10p::LNG1 L5) cotyledons. E. FWHM values scale with cotyledon anisotropy. Same n values as (B) and (D).

We measured SCD alignment in multiple independent transgenic lines for all three genotypes at 3 dpg and compared the FWHM values to those from Col-0 (Figure S6G). Increasing cotyledon elongation was sufficient to reorient the SCD field, with the highly elongated UBQ10p::LNG1 cotyledons showing the highest degree of stomatal alignment (Figure 6A-E, Figure S6D). This effect became more pronounced by 5 dpg as the UBQ10p::LNG1 cotyledons continued to lengthen at the expense of widening (Figure S6C,E). TMMp::LNG1 cotyledons showed an intermediate phenotype, with FWHM values between those from Col-0 and UBQ10p::LNG1 (Figure 6A-E). Interestingly, although SPCHp::LNG1 seedlings made slightly longer cotyledons, stomatal alignment did not differ significantly from Col-0, suggesting the existence of a threshold below while increasing elongation is insufficient to alter SCD orientation (Figure S6A,B).

While our ablation data demonstrated that directional cell expansion reorients SCDs, our LNG1 overexpression lines gave us the opportunity to investigate more specifically whether SCD orientation is driven by changes in GMCs or by a non-autonomous effect of expansion in the surrounding pavement cells. We quantified GMC morphology and found that neither GMC area nor the GMC aspect ratio were changed in any of our LNG1 lines (Figure S7A-C). In stark contrast, pavement cell expansion strongly reoriented along the proximodistal axis, and the magnitude of expansion scaled with both the strength of the promoter and the degree of SCD hyper-alignment (Figure S7D,F,G). These data indicate the global change in SCD orientation was not driven by increasing GMC anisotropy, but rather was due to a non-autonomous effect of directional cell expansion in surrounding pavement cells on the orientation of GMC long axes.

Our LNG1 overexpression lines demonstrated that hyper-alignment with the proximodistal axis is tunable; the greater the anisotropic pavement cell expansion along that axis, the higher the stomatal alignment. Finally, we tested whether shifting expansion laterally could reorient the SCD field away from the proximodistal axis. To accomplish this, we overexpressed *OVATE FAMILY PROTEIN 2* (*OFP2*) under the control of the UBQ10p (UBQ10p::OFP2). These lines exhibited the phenotypes seen in previously reported OFP overexpression lines(68–70), with notably squat cotyledons and true leaves that were readily distinguishable from those produced in Col-0 (Figure 6A). In agreement with our LNG1 data, changing the major direction of cotyledon expansion in UBQ10p::OFP2 was sufficient to reorient the stomatal field (Figure 6A-E). The altered SCD field in 3 dpg UBQ10p::OFP2 fanned outward along the margins while remaining aligned in the proximodistal direction near the midvein. The lateral reorientation became more pronounced over development, and the SCD distribution became bimodal by 5 dpg (Figure S6C,F). Finally, as in the LNG1 overexpression lines, GMC morphology was not altered in UBQ10p::OFP2 but pavement cell expansion was reoriented in the lateral direction (Figure S7). Therefore, the stomatal field is plastic in *Arabidopsis* and directly tunable by altering the major axis and magnitude of pavement cell expansion.

### SCD alignment is widespread across eudicots and is oriented along tissue axes

The mechanism of SCD orientation we found in *Arabidopsis* could explain stomatal alignment patterns across species without invoking unknown, conserved polarity regulators. Indeed, our data are consistent with previously published observations in monocots and would be sufficient to explain the hyper-alignment in those species. Therefore, we explored the possibility that cryptic SCD fields might be more widespread in eudicot leaves than previously appreciated by quantifying stomatal alignment in morphologically varied true leaves from seven additional eudicot species from different genera: *Carica papaya*, *Eschscholzia californica*, *Helianthus annuus*, *Lactuca sativa*, *Medicago sativa*, *Nicotiana benthamiana*, and *Solanum lycopersicum* (Figure 7A). We analyzed global SCD alignment in each species, focusing on midvein-adjacent regions, and compared the measurements to the FWHM scores from both *Arabidopsis* and maize. Because leaf size and stomatal number varied dramatically across the different species, we performed whole-leaf annotations for particularly informative cases.

**Figure 7:**
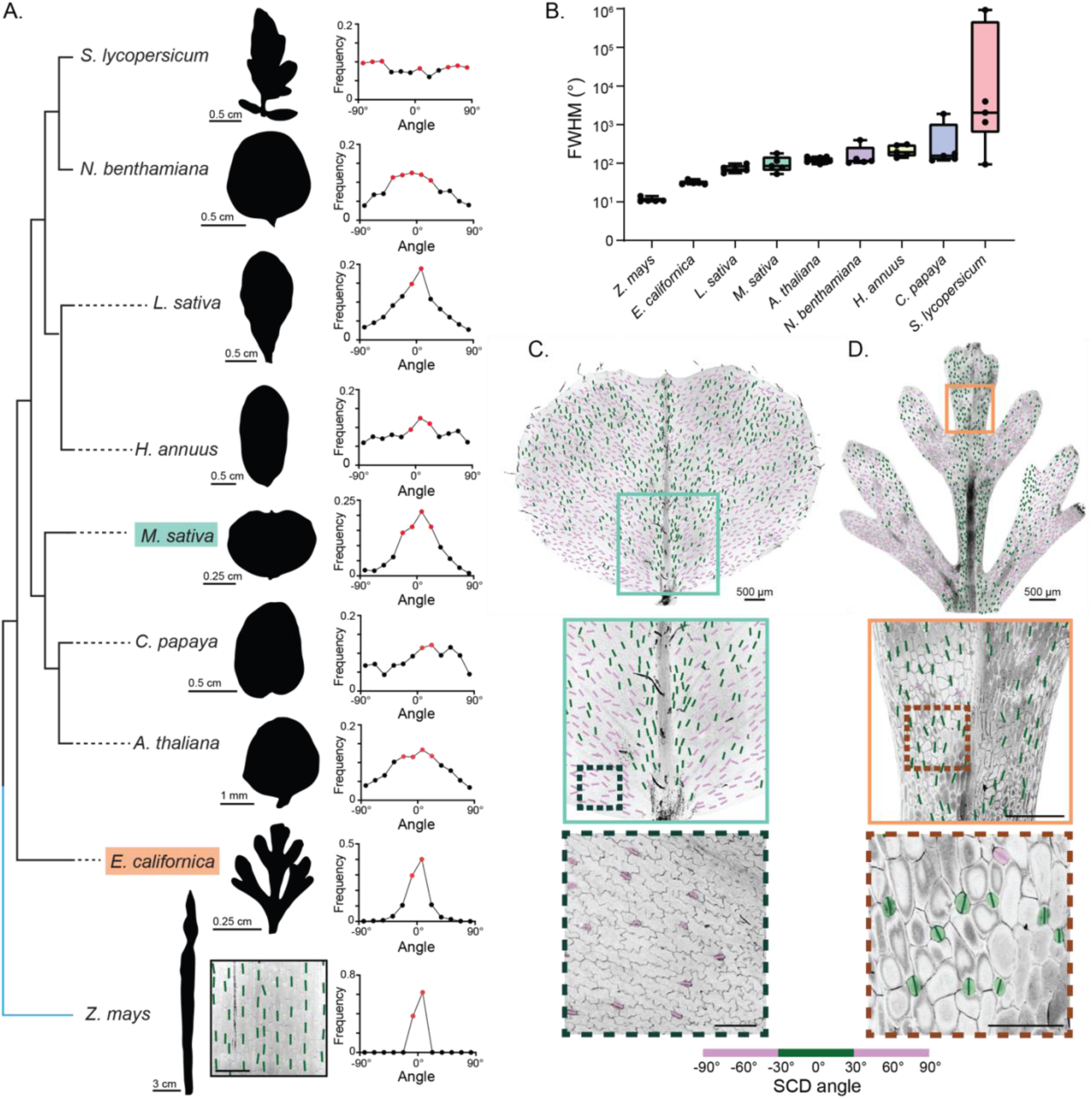
Oriented SCD fields are widespread across eudicots and align stomata to tissue axes. A. (Left) Representative images of true leaves that were used for SCD quantification with the phylogenetic relationships between the investigated species. (Right) Stomatal alignment relative to the proximodistal axis along the midvein for each species. Scale bars – 0.5 cm (*S. lycopersicum*, *N. benthamiana*, *L. sativa*, *H. annuus*, *C. papaya*), 0.25 cm (*M. sativa*, *E. californica*), 1 mm (*A. thaliana*), 3 cm (*Z. mays* whole leaf) and 300 μm (*Z. mays* zoom). B. FWHM values for indicated species. n = 5 true leaves each except *A. thaliana* (n=11). C. Representative image of a *M. sativa* true leaf with color-coded SCDs. Scale bars – 500 μm (top and middle images) and 100 μm (bottom image). D. Representative image of an *E. californica* true leaf with color-coded SCDs. Scale bars – 500 μm (top image), 250 μm (middle image) and 100 μm (bottom image).

Polarized SCD fields were found in all of the examined species, although both the magnitude and orientation of global alignment varied significantly (Figure 7B). *L. sativa*, *N. benthamiana*, *H. annuus* and *C. papaya* displayed patterns much like *Arabidopsis*, with divisions biased along the leaf’s proximodistal axis. SCD orientation near the midvein in *S. lycopersicum* true leaves was variable, with some of the samples showing high proximodistal alignment and others aligning in the mediolateral direction. Because of this variability, we checked near serrations at the leaf margin and found that stomata were aligned along the direction of outgrowth, similar to what we observed in *Arabidopsis* true leaf serrations.

Of the species we examined, we were the most intrigued by the SCD orientation fields seen in the elongated leaflets of *E. californica* and the heart-shaped leaves formed by *M. sativa* (Figure 7C,D) which mirrored key leaf shape features we saw in our UBQ10p::LNG1 and UBQ10p::OFP2 lines, respectively. Strikingly, stomata were hyper-aligned along the proximodistal axis of each leaflet in *E. californica* while stomata in the *M. sativa* true leaf were aligned along the proximodistal axis near the midvein but fanned outward toward the leaf margins nearer to the periphery. In both species, a closer look at cell morphology revealed that stomata were aligned with the direction of pavement cell expansion. Therefore, the stomatal patterns we found in other eudicots are consistent with the force-responsive SCD orientation model we identified in *Arabidopsis*.

## Discussion

In this work, we identified a new stomatal patterning mechanism that relies on orientation of the final symmetric cell division in the lineage. This regulation creates a polarized stomatal field in multiple organs, with guard cell orientation biased along global tissue axes. While stomatal alignment had been noted in some species for decades(24), it was generally considered to be restricted within certain genera and, importantly, the underlying mechanisms were unknown. Our finding that stomata are aligned in *Arabidopsis* leaves provided an ideal system to identify the responsible pathways.

Based on our analyses of ACD mutants (*basl*, *brx-q* and *tmm-1*), stomatal alignment does not share common regulators with those that control the one-cell spacing of stomata. Additionally, perturbation of auxin signaling, a key regulator of cell division in other tissues, did not reduce SCD alignment along tissue axes, leading us to consider alternative pathways capable of both controlling division orientation in individual GMCs and linking division orientation in a shared direction across the epidermis. Our results indicate that force-responsive mechanisms are responsible for controlling SCD orientation at both levels.

Using time-lapse microscopy, we found that GMCs divide along their long axes, which could be explained by 1) a cellular ruler that “measures” a cell’s dimensions or 2) a mechanics-based pathway that aligns divisions along the axis of greatest strain, which in wild-type cells will tend towards the longest axis. We found that SCDs no longer follow the GMC long axis in *KTN1* mutants even as GMC morphology remains unchanged, supporting a force-based mode of division orientation(58). How, then, is the division plane oriented along the axis of greatest strain? Our data indicate that the microtubule network is less anisotropic and less aligned with the GMC long axis in *ktn1-2*, consistent with a role for microtubule self-organization in division orientation. One of our more puzzling findings was that GMCs continue to divide along their long axes in *trm678*, suggesting that proper preprophase band establishment is dispensable for SCD orientation. This finding could be explained by a mechanism where interphase arrays influence division orientation independent of PPB establishment, which is supported by some recent work(71). Alternatively, ultrastructure studies have found that the cell walls are slightly thicker at the poles along the GMC long axis(72). Whether microtubule alignment along the long axis is responsible for this cell wall thickening and whether cell wall anisotropy is immediately upstream of division orientation will need to be tested in future studies.

By tracking cell and leaf expansion over time, we showed that GMC growth along the principal direction of leaf expansion primes the SCD field. Accordingly, mutants that alter leaf shape reorient SCD orientation. A key, outstanding question remains how the growth of individual cells is coordinated with growth of the developing organ. In plants, mutants that alter cell number or shape do not necessarily show changes in whole-organ morphology(73, 74). This suggests that, while changes in division or morphology at the cellular level are typically coordinated with development of the tissue, these processes can be uncoupled and tuned independently of each other. Further dissection of this via computational modeling could be useful to untangle the relative contributions of cell expansion, cell wall remodeling and cellular composition of the epidermis on this process.

The mechanics-based division orientation program outlined here provides a parsimonious explanation for the long-observed stomatal patterns seen in grasses and other plant species. Our survey of eudicots suggests that stomatal alignment is found more broadly than previously thought. Indeed, stomatal alignment in *Arabidopsis* is quite robust but has gone unnoticed until now, potentially because quantification of SCD alignment across the entire tissue had not been done. We suspect that revisiting other plant species using a more comprehensive analysis pipeline that quantifies stomatal orientation across larger areas will uncover significantly more species with aligned stomata. While the model we put forth here is sufficient to explain the diversity of stomatal alignments we found, conclusive proof of conservation will require both the identification of the relevant molecular regulators and the ability to perturb those regulators in non-model plant species.

Proper division orientation is one of the pillars of morphogenesis. In some cases, such as in the dome-shaped shoot apical meristem, oriented divisions control overall organ shape^56^. The oriented SCDs we described here are notably different in this regard. We have not found any evidence to suggest that stomatal lineage SCDs impact overall organ (e.g. cotyledon, true leaf) morphology. Instead, when SCDs are misoriented, as they are in *ktn1-2*, they create aberrantly shaped stomata with small pores. Interestingly, malformed stomata were observed in a previous study(75), and our data identify the underlying defect. We found that KTN1 is not required for symmetric division *per se* but, rather, is necessary for robust divisions along the GMC long axis. This reveals an unexpected function for SCD orientation in determining stomatal morphology and indicates that the characteristic kidney-bean shape of *Arabidopsis* GCs is not solely a product of post-division cell expansion. Why misoriented SCDs lead to GC morphology and pore defects remains to be determined. At the moment, we favor a model where oriented SCDs are required to coordinate the position of cell wall thickenings with the division plane, but future exploration of cell wall deposition in misoriented *ktn1-2* SCDs will be required to clarify this point. Finally, how the increased randomization of SCD orientation in *ktn1-2* affects stomatal kinetics remains to be seen. It is tempting to speculate that global stomatal alignment could influence overall stomatal physiology by impacting pore opening and closing efficiency. By identifying additional genes required for SCD orientation in the leaf epidermis, we can begin to test how the polarized stomatal field influences gas exchange to regulate photosynthesis and plant growth.

## Acknowledgements

We are grateful to Dr. Mark Estelle, Dr. Alisa Huffaker, Dr. Alexandra Dickinson and Dr. Dominique Bergmann for seeds. We thank the Muroyama lab members for their input and helpful suggestions. We want to thank Ivan Jaramillo, Aranza Martinez Leija, Lizbeth Rodriguez and Daniella Van Maele for technical assistance. We also remember and thank Nathaniel Takatsuno for productive discussions, technical assistance, and his joyful spirit. We dedicate this work to him. This work was supported by funds from UCSD, R35 GM150466, and a Hellman Fellowship, all to A.M.

## Author contributions

Conceptualization – K.S.H. and A.M. Software – K.S.H. Investigation – K.S.H., B.Y.L, J.H.G., M.E.G., A.C. and A.M. Formal Analysis – K.S.H., B.Y.L, J.H.G., M.E.G., A.C. and A.M. Writing – K.S.H. and A.M. Visualization – KS.H. and A.M., Supervision – A.M. Funding Acquisition – A.M.

## Supplemental information

Document S1. Figures S1-S7.

## Methods

### Experimental Model

All *Arabidopsis thaliana* lines generated and used in this study were in the Col-0 background. ML1p::mCherry-RCI2A(76), BRXL2p::BRXL2-YFP ML1p::mCherry-RCI2A(30), BASLp::YFP-BASL 35Sp::PIP2A-RFP(30), BRXL2p::BRXL2-YFP TMMp::mCherry-TUA5(55), *basl-2* (WiscDsLox264F02)(9) 35Sp::PIP2A-RFP, *brx-q*(*30*), *tmm-1*(*35*), *flp-1 myb88*(*36*), *trm678*(*56*), *ktn1-2* (SAIL_343_D12)(77) were published previously. *kan1-11 kan2-5*(*61*) (AT5G16560 and AT1G32240, CS67888), *rot3-1*(*62*) (AT4G36380, CS3727), and *scf-9*(*44*) (AT5G13300, SALK_069166) were acquired from the Arabidopsis Biological Resource Center at Ohio State University and the ML1p::mCherry-RIC2A reporter was introduced by crossing. *opl1-1 opl2-3 opl3-1 opl4-1*(*34*) (*opl-quad*) and *fama-1*(*37*) were previously published and kindly provided by Dr. Dominique Bergmann (Stanford/HHMI). *tir1-1 afb2-3 afb3-4* was published previously(43) and kindly provided by Dr. Mark Estelle (UCSD). *kan1-11 kan2-5* and *rot3-1* were confirmed by PCR. *kan1-11* was genotyped by first amplifying the mutation-harboring locus via PCR using CTCTCCAGTTTGTCATCTG and ACCACTCAACTTTAGGGTTC and sequencing using CATCTGTAATTCTGTATC for a missense mutation in the 3^rd^ exon (G1801A, R272Q). *kan2-5* was amplified with GGTTCATCATCTGTGGAAACCG and CGAGTAATTCAACGGCGTGAAC and sequenced with GTAAACCATCATCGACATGGfor the presence of the nonsense mutation (C268A). *rot3-1* was genotyped using PCR followed by gel electrophoresis using CGAGACAAAACGGCCTAAGC and AAGTTTAGGGTTTCTCCGATCACC to validate the deletion. *scf-9* and *tir1-1 afb2-3 afb3-4* were confirmed by mounting analyzed seedlings in Hoyer’s solution and confirming the previously described vein morphology defects.

*Arabidopsis* seeds were sterilized in 20% bleach for 10 min, followed by 3 rinses with Milli-Q water. Seeds were then sown on ½ MS + 0.5% sucrose plates and stratified in the dark at 4°C for at least 2 days. The day a plate was moved into the growth chamber was counted as 0 dpg. The growth chamber (Percival, Model CU41L5) was set to long-day conditions (16 h light, 8 h dark) at 22°C with SciWhite^TM^ LED lighting at 650 lux. The IAA experiments and Col-0 needle ablations were performed in the same Percival with identical light conditions but set to 25°C.

Tobacco (*N. benthamiana*), tomato (*S. lycopersicum* cv. M82) and maize (*Z. mays*, B73) were generously provided by Dr. Alexandra Jazz Dickinson (UCSD). Alfalfa (*M. sativa*, accession W6 2502) was a kind gift from Dr. Alisa Huffaker (UCSD). Lettuce (*L. sativa* cv. Parris Island cos), California poppy (*E. californica* cv. aurantiaca orange), and sunflower (*H. annuus* cv. Russian mammoth) were purchased from Home Depot (San Diego, CA). Papaya (*C. papaya*, ‘Mexican’) seeds were harvested from a papaya fruit bought at a local grocery store.

Tomato seeds were sterilized via incubation in 70% ethanol for 5 min followed by incubation in 40% bleach for 30 min and finally 3 rinses with Milli-Q water. Seeds were sown on ½ MS + 0.5% sucrose plates with the root end of the seed facing downward on an upright plate. Seeds were stratified in the dark at 4°C for 3 days before being grown vertically in the growth chamber under the same conditions as *Arabidopsis*. Once cotyledons emerged and roots were sufficiently established, seedlings were transferred to soil.

Sunflower, lettuce, poppy, and tobacco seeds were sown on moist soil. Individual tobacco seedlings were split into separate pots ∼10 days after sowing. Alfalfa seeds were sterilized and plated on ½ MS plates following the *Arabidopsis* protocol and moved to soil once germinated.

Maize seeds were prepared by first incubating in DI water for 4-6 h. Next, tip caps were removed using a razor blade, taking care not to damage the embryo and endosperm. Seeds were then incubated in warm water (55-57°C) for 5 min, and then rinsed once with room temperature water. Next, seeds were incubated in 20% bleach for 20 min, and the tube was shaken every few minutes. In a laminar flow hood, seeds were rinsed 5-7 times with autoclaved DI water. Lastly, seeds were sandwiched between moist, autoclaved paper towels, set in a square plate with all cut ends pointing the same direction, and the plate was sealed with Parafilm. Plates were grown vertically in a growth room set to 29°C with long-day conditions (16 h light, 8 h dark). Once seeds had germinated, they were moved to soil and continued in the growth room.

Papaya seeds were prepared by removing the aril (the germination-inhibiting coat around the seed) with tweezers. Seeds were sterilized in 20% bleach for 15 min, then plated in paper towels, following the maize protocol. Seeds were germinated at 29°C and then transferred to soil and grown at 22°C.

## Materials and Methods

### Generation of plasmids for plant transformation

TMMp::LNG1, SPCHp::LNG1, UBQ10p::LNG1, and UBQ10p::OFP2 were all generated using Gateway cloning. LNG1 (AT5G15580) was amplified from *Arabidopsis* genomic DNA using CACCCAACAACCTTCTGAGGCCAGAG and GGGGTTCAGAGAACCAAGAAACC and ligated into pENTR D-TOPO. OFP2 (AT2G30400) was amplified from *Arabidopsis* genomic DNA using CACCATGGGGAATTACAAGTTCAG and TTACTTTGTTTTTGTAAGTTG and ligated into pENTR D-TOPO. The TMM and SPCH promoter sequences were in pDONR P4-P1R(78) and the UBQ10 promoter sequence was in pENTR 5’ TOPO. Vectors were assembled in R4pGWB601(79). All constructs were validated by sequencing.

### Generation and origin of transgenic lines

To generate all OFP2- and LNG1-overexpressing lines, plants containing the plasma membrane reporter ML1p::mCherry-RCI2A were transformed using the agrobacterium-mediated floral dip method(80) in strain GV3101. Transformants were selected on ½ MS plates with 150 μM PPT and no sucrose. Mutant lines *kan1-11 kan2-5* (AT5G16560 and AT1G32240, CS67888) and *rot3-1* (AT4G36380, CS3727) were obtained from the ABRC at Ohio State University. The plasma membrane reporter construct ML1p::mCherry-RCI2A was introgressed by crossing.

### Confocal microscopy

All confocal microscopy, with the exception of the images of microtubule organization in GMCs and guard cells (Figure 3C and Figure S3J), was done on a Leica Stellaris 5 with HyD detectors using the 10x/0.40NA or 20x/0.75NA objectives. For *Arabidopsis* lines without genetically encoded plasma membrane markers, cell walls were stained by incubating samples in propidium iodide (PI, 10 μg/ml) for 5 min. For non-*Arabidopsis* species, cell walls were stained by incubating in PI (10-200 μg/ml, higher concentration used for larger samples) for 30 min-7 hr or with FM4-64 (20-40 μM) for 2-7 h (alfalfa, lettuce, maize, sunflower, tomato) depending on the time required to successfully stain the cell walls. PI and YFP reporters were excited by a 514 nm laser and their emissions were detected between 520-650 nm. FM4-64 and mCherry reporters were excited by a 561 nm laser and their emissions were detected between 565-650 nm. Time-lapse imaging for determining the GMC’s long axis prior to division was acquired with 90 min time intervals and a duration of 13-15 h using the 20x objective at 2x zoom. Whole seedlings were mounted on a coverslip and covered with a 0.15 mm HybriWell chamber (Grace Bio-Labs, HBW75) connected to a pump (SyringePump.com, NE-300 Just Infusion™ Syringe Pump) flowing ½ MS liquid media at 0.04 ml/min. For manual time course imaging (images acquired 12 or 24 h apart), whole seedlings were gently placed on and removed from slides without damaging the root and were placed back on plates in the growth chamber between imaging sessions. Whole leaf images (Figure 7) were acquired on a Samsung Galaxy phone camera.

For mounting sepals, flowers were dissected off the plant and mounted per the protocol published in He et al.(81) The cut stem was attached to a glass slide with tape, so the flower was lying on its side with one sepal facing up. After adding a drop of water, a coverslip was taped on top to flatten the sample.

Images of microtubules were obtained on a Nikon Eclipse Ti2-E microscope with a CSU-X1 spinning disk, a Prime 95B sCMOS camera and the Plan Apo ʎ 60x NA 1.40 oil objective.

### Image analysis

Most analyses were performed on max projections created from z-stacks using Fiji. SCD orientation relative to the leaf’s proximodistal axis was measured in Fiji by manually annotating the cell wall separating paired guard cells using the “Straight Line” tool. For analysis of synchronous and asynchronous SCDs of stomatal pairs in *basl-2*, pairs where the stomata were roughly the same size were considered synchronous, and otherwise they were classified as asynchronous. GMCs were measured by manually tracing the cell outline with the “Polygon tool” and fitting an ellipse in Fiji to find the long axis, aspect ratio, and area. When comparing GMC morphology with corresponding division angle, the GMC was measured in the time frame immediately preceding the time frame in which the division first appears (i.e., 90 min before for Col-0 and *ktn1-2* movies, 60 min for *trm678* movies). The difference between the GMC long axis and SCD angle was always calculated as the acute angle; to accommodate this, the range of the angles was extended up to 120°.

To measure whole cotyledon anisotropy, the outline of each sample (with the petiole trimmed from the image) was first extracted using Fiji’s “Analyze Particles” tool. For most samples, anisotropy was defined as the AR calculated by fitting an ellipse to this outline. For samples that were visually wider than they were long, anisotropy was defined as the inverse of the AR. Finally, for samples where the calculated longest axis of the fit ellipse did not align with either the proximodistal or lateral axis, the length of the sample along these two axes was manually measured using Fiji’s “Straight Line” tool, and the anisotropy was defined as the proximodistal length divided by the lateral length.

Quantification of microtubule anisotropy and major axis was performed using the FibrilTool plugin for Fiji(82). Pore size was analyzed by manually tracing the pore aperture using the “Polygon” tool in Fiji from maximum projections of the ML1p::mCherry-RCI2A reporter.

### SCD visualization, simulation, and averaging

The graphs with SCDs color-coded by angle were generated in MATLAB using annotation coordinates obtained in Fiji. For visualization purposes, the length of each SCD line was set to a user-defined length.

MATLAB was also used to find the average SCD angle by position on the leaf. For each sample, a 9×10 grid was fit to the cotyledon’s outline. For each grid space, the angles of SCDs inside its bounds were stored with those of the same grid space on all the other samples. SCDs straddling two grid spaces were counted in both spaces. These aggregated angles were then averaged, yielding the average orientation of all samples within each grid space. The standard deviation of angles within each grid space was also calculated. For plotting the averages, the thickness of each line was weighted by the number of SCDs that fell within that grid space.

Simulations of cotyledons with randomly oriented SCDs were generated in MATLAB. For each sample, an ellipse of random length and width was generated from the normal distribution of 3 dpg cotyledons lengths and aspect ratios, yielding the simulated cotyledon outline. 150 (the average stomatal number at 3 dpg) randomly oriented SCDs were generated within the ellipse bounds.

### FWHM global alignment analysis

Full width at half maximum (FWHM) was calculated in MATLAB. A one-term Gaussian model was fit to the histogram of SCD angles, yielding the following model: f(x) = a*e^-((x-b)/c)^2^. FWHM was then calculated using the following equation: FWHM = 2*sqrt(log(2))*c.

### AFT local alignment analysis

Alignment by Fourier Transform (AFT) was performed using the Matlab package described in Marcotti et al.(29) This analysis quantifies the alignment of fibrils or lines in an image, outputting a score from 0 to 1, where 0 reflects random alignment and 1 indicates complete alignment between neighbors. An image is divided into a grid of tiles of a user-defined ‘Neighborhood size’ and the average orientation within each tile is compared to a user-defined number of surrounding grid layers (‘Neighborhood radius’). The input images were generated in Fiji by flattening the manually annotated ROIs of the shared guard cell walls onto a blank image of the same dimensions as the original confocal image. When running ‘AFT_batch,’ the ‘Neighborhood size’ was set to the average area that contained an average of three stomata for a given genotype, developmental stage, or species. The ‘Window overlap’ was kept at the default 50% and the ‘Neighborhood radius’ was adjusted as needed. A larger ‘Neighborhood radius’ compares the local average SCD angle to an increasingly broader area of neighboring local averages. For pairwise comparisons in this manuscript, ‘Neighborhood radius’ was set to 3.

### Hormone treatments

For the IAA experiments, seeds were sown on ½ MS media with 0.5% sucrose containing either 500 nM or 1 µM IAA (Thermo Scientific Chemicals, AC122160100) or EtOH and grown at 25°C. Efficacy of the treatment was confirmed by visual phenotyping of seedlings, which displayed delayed and stunted growth. SCD phenotypes were evaluated at 3 dpg. For the NPA experiments, seeds were sown on ½ MS media with 0.5% sucrose containing either 10 µM NPA (Tokyo Chemical Industry, N0067) or DMSO and grown at 25°C.

### Hierarchical cluster analysis of cell growth data

Cotyledons of intact, 2 dpg seedlings were imaged at 12 h intervals, for a total of three images per sample. Between imaging sessions, seedlings were gently placed back on MS plates and grown in the growth chamber under our standard growth conditions. GMC long axis and SCD orientation relative to the proximodistal axis were first measured in Fiji. Then, for each GMC and SCD, the orientation relative to the average SCD angle at its position on the leaf at 3 dpg was calculated in MATLAB: first, each GMC and SCD was assigned to a grid space (i.e., position on the leaf), using the same method as was used for assigning SCDs when calculating the average, composite cotyledons (see Image Analysis); next, the difference between the measured GMC/SCD angle and the average 3 dpg SCD angle at the corresponding leaf position was calculated, yielding the GMC’s/SCD’s orientation relative to that average. Because the average SCD orientation varies by position on the leaf (e.g., those near the base tend to align along the margin), analyzing the data in this way reveals how well GMCs are oriented towards their position-specific, expected SCD orientation. To perform hierarchical clustering, cells were first classified into three groups based on the alignment of the GMC long axis at time = 0 hrs (2 dpg) with the average 3 dpg SCD angle at that position: aligned (0-30 degrees), mid-aligned (30-60 degrees), and unaligned (60-90 degrees). For each group, the Euclidean distance metric of each cell’s angle over time was calculated using the dist() function in R. The hclust() function was then run on the Euclidean distance metric for each group, using the UPGMA agglomeration method. Hierarchical clustering groups together similar trajectories without inherently defining the number of clusters, requiring the user to define the number of clusters after the fact. At 24 h, SCDs will fall within one of the three ranges defined initially as aligned, mid-aligned, or unaligned. Based on the prediction that each group might naturally cluster into those three ranges for the 24 h SCD angle, the number of clusters was set to three.

### Needle ablation

Small regions of cells, between 2724 µm^2^ and 19,306 µm^2^, on the abaxial epidermis of 3 dpg cotyledons were ablated using a 0.25 mm tungsten needle (Roboz Surgical Instruments, RS-6064). Identically sized, non-damaged regions on the contralateral side of the same leaf were used as controls. Immediately after ablation, intact seedlings were mounted and cotyledons were imaged to capture the orientation of existing GMCs and SCDs at 0 h. Seedlings were carefully placed back on MS plates without damage to the roots and grown 24 h in the growth chamber under normal conditions. After 24 h, cotyledons were imaged again, to capture the orientation of SCDs occurring within the previous day.

The orientation of GMCs and SCDs were measured relative to the cotyledon’s proximal distal axis using Fiji. A MATLAB script was then used to generate the “radial angle,” the angle between the SCD’s angle relative to the leaf axis and the angle of a line from the center of the SCD to the ablation or control center. For 0 h, the ablation/control center was determined with MATLAB by fitting an ellipse to the region’s outline. For 24 h, the center was manually measured in Fiji by tracking cell landmarks between 0 and 24 h. To visualize cell growth around the ablation site, the layer of cells immediately surrounding the ablation at 0 h and 24 h was traced in Illustrator. To account for overall expansion of the leaf, the 24 h trace was scaled down to fit within the same size bounding circle that fit around the 0 h trace, and the traces were overlaid by aligning the coordinates of the ablation center at both time points.

### Quantification and statistical analysis

Gene sequences were obtained from The Arabidopsis Information Resource (TAIR). Statistical analysis and graph creation was done using GraphPad Prism (version 10.2.3). The means of two groups were compared using an unpaired t test. For comparing multiple groups, one-way ANOVA was used. If there was a significant difference between any of the means (p-value <0.05), a Dunnett test was performed to compare the mean of each group with the control group’s mean (Col-0). Sample sizes and p-values are reported in figure legends. Error bars represent the standard deviation. Hierarchical clustering was performed in R and Rstudio (version 2023.06.1+524.pro1) using the tidyverse and gghighlight packages. Image analysis was performed with Fiji/ImageJ (version 2.14.0/1.54f). Analyses done in MATLAB utilized release R2024a. Cell tracing around ablation sites was performed in Adobe Illustrator (Beta) (version 29.8).

---

**Figure S1:**
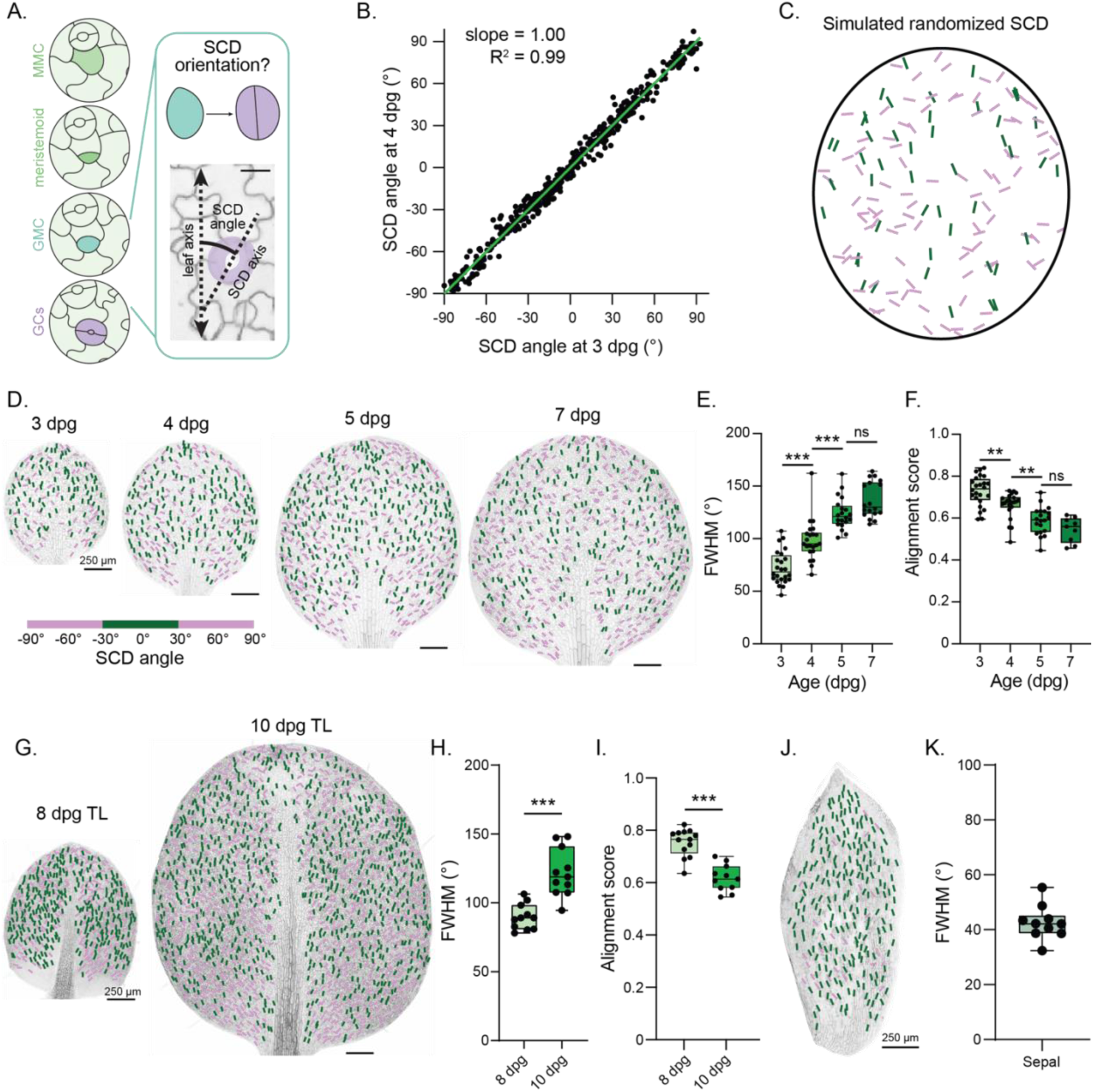
The polarized SCD field is developmentally regulated and is found across other stomata-containing tissues. A. (Left) Schematic of the *Arabidopsis* stomatal lineage. (Right) Method to measure stomatal alignment to the proximodistal axis. Scale bar – 20μm. B. Comparison of SCD angles of paired stomata tracked from 3 dpg to 4 dpg. n = 490 cells. C. Example of a simulated cotyledon with randomized, color-coded SCDs. D. Representative images of ML1p::mCherry-RCI2A-expressing cotyledons with color-coded SCDs at the indicated developmental stages. Scale bars – 250 μm. E. FWHM values across cotyledon development. n = 24 (3 dpg), 22 (4 dpg), 20 (5 dpg), 18 (7 dpg) cotyledons. Note that the 3 dpg data are the same as those in Figure 1E. F. Alignment scores across cotyledon development. Same n values as (E) except for 7 dpg (n = 9). G. Representative images of 8 and 10 dpg true leaves (TLs) with color-coded SCDs. Scale bars – 250 μm. H. FWHM values for 8 and 10 dpg true leaves. Note that the 10 dpg data is the same as those shown in Figure 7B. n = 11 true leaves each. I. Alignment scores for the 8 and 10 dpg true leaves in (H). n = 13 (8 dpg) and 11 (10 dpg) cotyledons. J. Representative image of a sepal with color-coded SCDs. Scale bar – 250 μm. K. FWHM values for sepals. n = 10. ns – not significant, ** - p<0.01, *** - p<0.001

**Figure S2:**
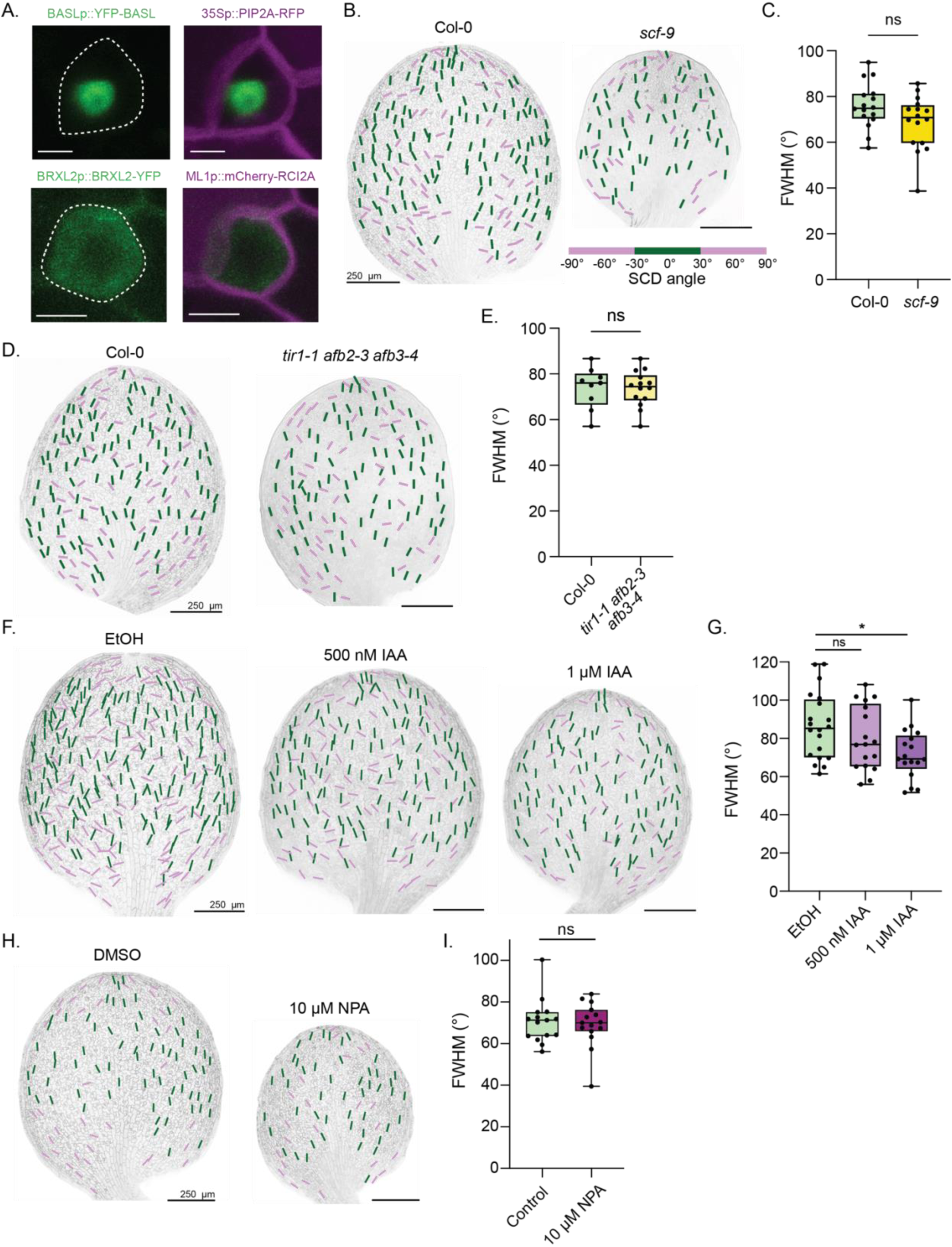
ACD polar proteins, vein patterning, and auxin do not control the orientation of the SCD field. A. Representative images of the BASLp::YFP-BASL (top) and BRXL2p::BRXL2-YFP (bottom) reporters in GMCs. Scale bars – 5 μm. B. Representative images of 3 dpg Col-0 and *scf-9* cotyledons with color-coded SCDs. Scale bars – 250 μm. C. FWHM values for Col-0 and *scf-9*. n = 16 cotyledons each. D. Representative images of 3 dpg Col-0 and *tir1-1 afb2-3 afb3-4* cotyledons with color-coded SCDs. Scale bars – 250 μm.

E. FWHM values for Col-0 and *tir1-1 afb2-3 afb3-4*. n = 9 (Col-0) and 14 (*tir1-1 afb2-3 afb3-4*) cotyledons.

F. Representative images of 3 dpg cotyledons from seedlings grown on plates with the indicated pharmacological treatments (EtOH as control, 500 nM IAA, and 1 μM IAA). Scale bars – 250μm.

G. FWHM values for 3 dpg cotyledons from seedlings grown on plates with the indicated pharmacological treatments. n = 20 (EtOH) and 17 (500 nM IAA and 1 μM IAA) cotyledons.

H. Representative images of 3 dpg cotyledons from seedlings grown on plates with the indicated pharmacological treatments (DMSO as control and 10 μM NPA). Scale bars – 250μm.

I. FWHM values for 3 dpg cotyledons from seedlings grown on plates with the indicated pharmacological treatments. n = 15 cotyledons each.

ns – not significant, * - p<0.05

**Figure S3:**
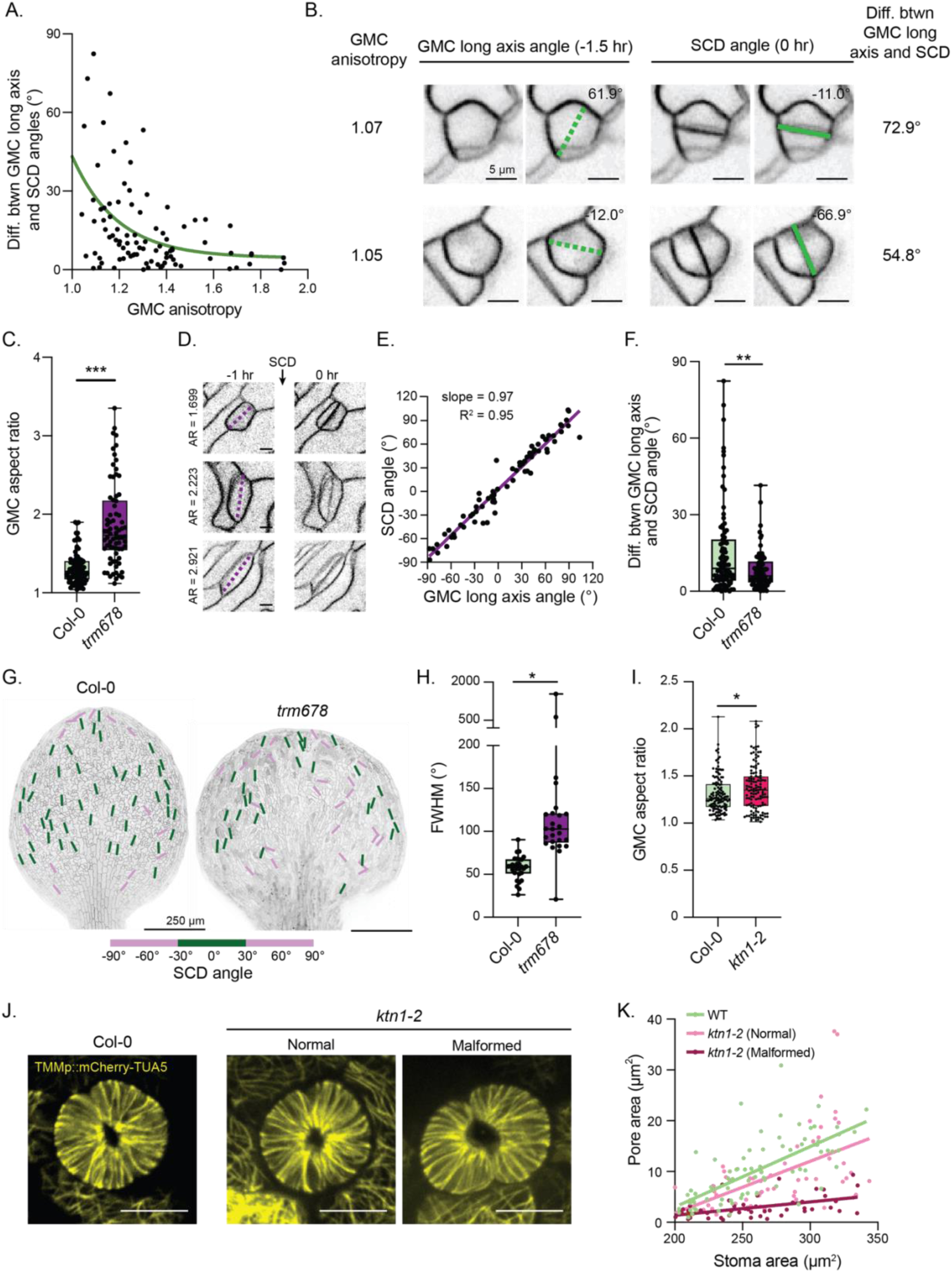
Additional characterization of the relationship between GMC morphology, SCD orientation, and stomatal pore formation. A. The difference between the GMC long axis angle and SCD angle as a function of GMC aspect ratio (green line – non-linear fit (one-phase decay)). n = 91 cells. B. Representative images of nearly isotropic GMCs where the SCD (right, solid green line) deviates significantly from the GMC long axis (left, dashed green line). Scale bars – 5 μm. C. Quantification of GMC aspect ratio in Col-0 and *trm678* GMCs. n = 91 (Col-0) and 74 (*trm678*) cells. D. Representative images of SCDs in *trm678* GMCs. The dashed purple line indicates the GMC long axis immediately before mitosis. Scale bars – 5 μm. E. Comparison of the angle of the GMC long axis immediately before mitosis to the associated SCD angle in *trm678* (purple line – linear fit). n = 74 cells. F. Differences between the GMC long axis angle and SCD angle in Col-0 and *trm678*. Note that the data for Col-0 are the same as those in Figure 3H. n = 91 (Col-0) and 74 (*trm678*) cells. G. Representative images of 3 dpg Col-0 and *trm678* cotyledons with color-coded SCDs. Scale bars – 250μm. H. FWHM values for Col-0 and *trm678*. n = 29 (Col-0) and 25 (*trm678*) cotyledons. I. Quantification of GMC aspect ratio in Col-0 and *ktn1-2* GMCs. n = 125 (Col-0) and 124 (*ktn1-2*) GMCs. J. Microtubule organization (TMMp::mCherry-TUA5) in Col-0 and normal and malformed *ktn1-2* stomata. Scale bars – 10 μm. K. Comparison of the stoma and pore areas in Col-0 and *ktn1-2*. n = 76 (Col-0), 57 (normal, *ktn1-2*), and 50 (malformed, *ktn1-2*) pores. * - p<0.05, ** - p<0.01, *** - p<0.001

**Figure S4:**
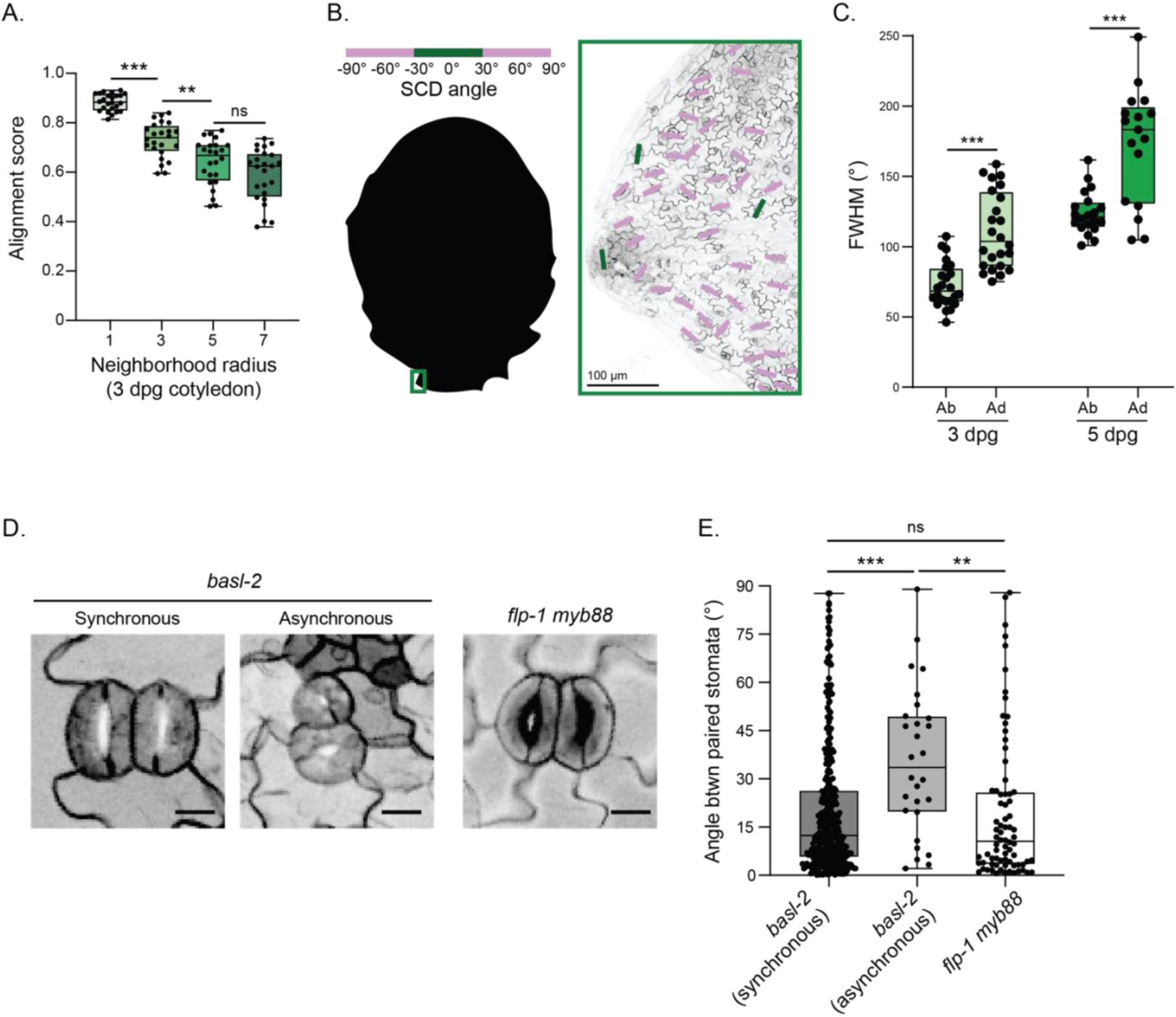
Additional characterization of Col-0 and stomatal cluster mutants implicates cell expansion as an important driver of SCD orientation. A. Alignment scores by varying neighborhood sizes in 3 dpg cotyledons. n = 24 cotyledons. B. Example image of an SCD field oriented toward the tip of a developing serration of an *Arabidopsis* true leaf. Scale bar – 100 μm. C. FHWM values for abaxial and adaxial SCDs at both 3 dpg and 5 dpg. n = 24 (3 dpg Ab), 24 (3 dpg Ad), 20 (5 dpg Ab) and 17 (5 dpg Ad) cotyledons. Note that the 3 dpg abaxial data are the same as those shown in Figure 1E, and the 5 dpg abaxial data are the same as those shown in Figure S1E. D. Representative images of stomatal pairs in *basl-2* and *flp myb88*. Scale bars – 10 μm. E. Differences between the orientation of paired stomata in *basl-2* and *flp myb88*. n = 352 (synchronous, *basl-2*), 28 (asynchronous, *basl-2*) and 76 (*flp myb88*) stomatal pairs. ns – not significant, ** - p<0.01, *** - p<0.001

**Figure S5:**
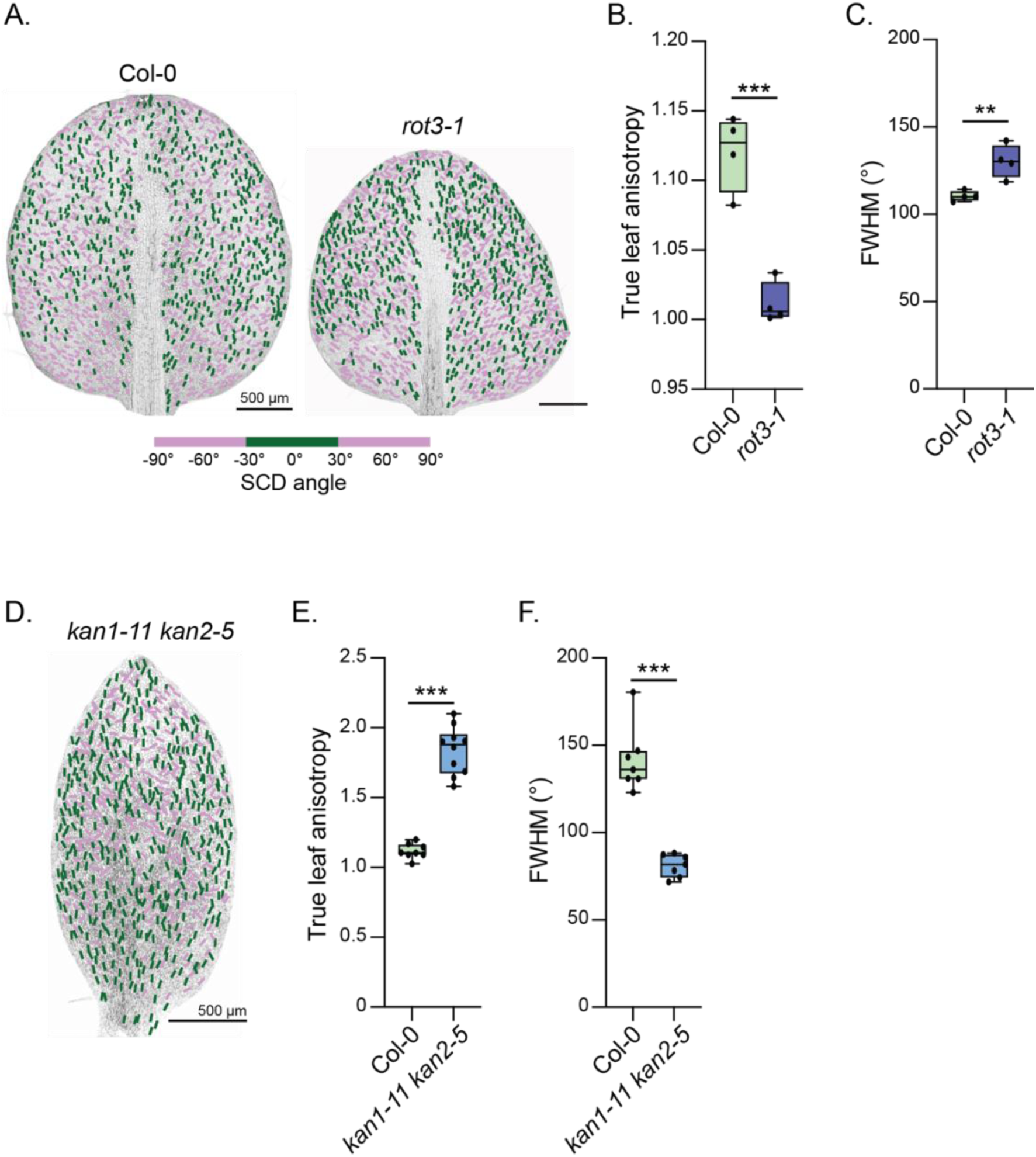
Mutations that affect leaf shape also reorient the polarized SCD field. A. Representative images of 10 dpg ML1p::mCherry-RCI2A-expressing Col-0 and *rot3-1* true leaves with color-coded SCDs. Scale bars – 500 μm. B. Anisotropy of 10 dpg Col-0 and *rot3-1* true leaves. n = 4 true leaves each. C. FWHM values for Col-0 and *rot3-1* true leaves in (B). n = 4 true leaves each. D. Representative image of a 10 dpg ML1p::mCherry-RCI2A-expressing *kan1-11 kan2-5* true leaf with color-coded SCDs. Scale bar – 500 μm. E. Anisotropy of 10 dpg Col-0 and *kan1-11 kan2-5* true leaves. n = 7 true leaves each. F. FWHM values for Col-0 and *kan1-11 kan2-5* true leaves in (E). n = 7 true leaves each. ** - p<0.01, *** - p<0.001

**Figure S6:**
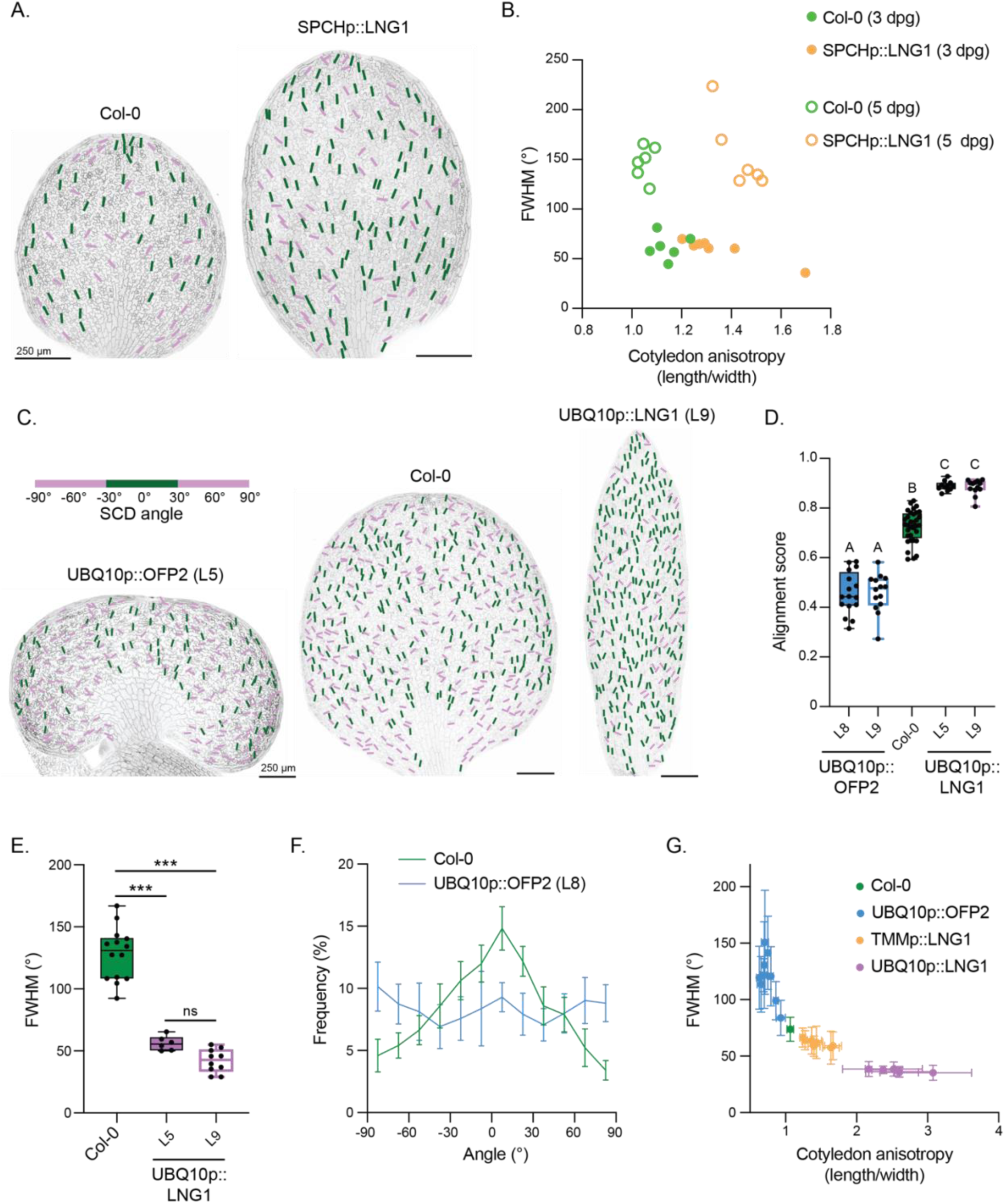
Additional characterization of LNG1 and OFP2 overexpression lines. A. Representative images of 3 dpg Col-0 and SPCHp::LNG1 cotyledons. Scale bars – 250 μm. B. Comparison of FWHM values and the associated cotyledon anisotropy for 3 dpg and 5 dpg Col-0 and SPCHp::LNG1 cotyledons. n = 6 cotyledons each except for 3 dpg SPCHp::LNG1 (n=7). C. Representative images of 5 dpg Col-0, UBQ10p::OFP2, and UBQ10p::LNG1 cotyledons. Scale bars – 250 μm. D. Alignment scores for the indicated genotypes at 3 dpg. n = 22 (Col-0), 16 (UBQ10p::OFP2 L8), 14 (UBQ10p::OFP2 L9), 11 (UBQ10p::LNG1 L5) and 13 (UBQ10p::LNG1 L9) cotyledons. E. FWHM values for the indicated genotypes at 5 dpg. n = 14 (Col-0), 6 (UBQ10p::LNG1 L5) and 10 (UBQ10p::LNG1 L9) cotyledons. F. Frequency distributions for SCD angles in the indicated genotypes at 5 dpg. n = 10 seedlings each. G. Comparison of FWHM values and the associated cotyledon anisotropy, where each point represents the mean value for independent transgenic lines of the indicated genotypes at 3 dpg. n = 9 (UBQ10p::OFP2), 7 (TMMp::LNG1), and 6 (UBQ10p::LNG1) independent transgenic lines. 8-18 cotyledons were analyzed for each line, with the exception of the paired Col-0 data (n = 55 cotyledons). ns – not significant, *** - p<0.001

**Figure S7:**
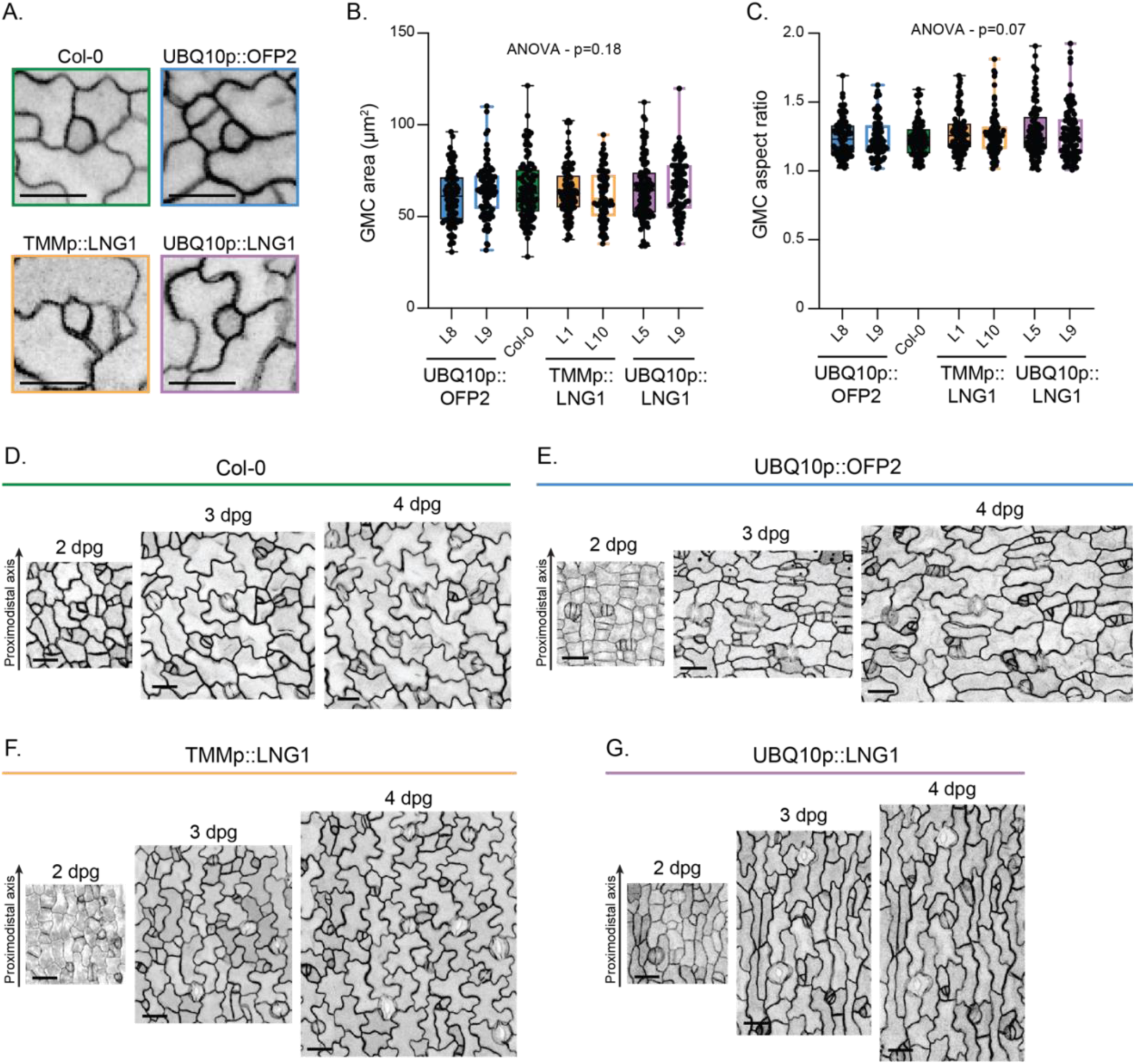
Additional characterization of pavement cell growth and GMC morphology in LNG1 and OFP2 overexpression lines highlights non-cell autonomous role of cell expansion on SCD orientation. A. Representative images of GMC morphology in the indicated genotypes (Col-0, UBQ10p::OFP2, TMMp::LNG1, and UBQ10p::LNG1). Scale bars – 20 μm. B. GMC area in 3 dpg cotyledons of the indicated genotypes. n = 98 (Col-0), 98 (UBQ10p::OFP2 L8), 97 (UBQ10p::OFP2 L9), 98 (TMMp::LNG1 L1), 90 (TMMp::LNG1 L10), 102 (UBQ10p::LNG1 L5) and 96 (UBQ10p::LNG1 L9) GMCs. C. GMC aspect ratio in 3 dpg cotyledons of the indicated genotypes (Col-0, UBQ10p::OFP2, TMMp::LNG1, and UBQ10p::LNG1). Same n values as (B). D-G. Representative image series of directional pavement cell expansion in the indicated genotypes (Col-0, UBQ10p::OFP2, TMMp::LNG1, and UBQ10p::LNG1). Scale bars – 25 μm.

